# The superior salinity tolerance of wheat cultivar Shanrong No. 3 cannot be attributed to elevated Ta-sro1 poly(ADP-ribose) polymerase activity

**DOI:** 10.1101/2021.10.20.465099

**Authors:** Sarah Vogt, Karla Feijs, Sebastian Hosch, Raffaella De Masi, Ruth Lintermann, Bernhard Loll, Lennart Wirthmueller

## Abstract

Saline soils limit the production of important staple crops such as wheat, particularly in arid and semiarid regions. Salt tolerance is a multi-gene trait and this complicates breeding of wheat varieties that deliver high yields under saline soil conditions. Notably, the elevated salinity tolerance of wheat cultivar Shanrong No. 3 (SR3) has been linked to a specific proteoform of the wheat SIMILAR TO RCD1 ONE (SRO1) protein that was created in an asymmetric genome hybridization with tall wheat grass. The two amino acid polymorphisms of the Ta-sro1 proteoform enhance the poly(ADP-ribose) polymerase (PARP) activity of the protein suggesting that altered poly-ADP-ribosylation of unknown substrate proteins or nucleic acids underlie the elevated salinity tolerance of cultivar SR3. To elucidate the molecular basis for the elevated PARP activity of the Ta-sro1 proteoform we solved a crystal structure of the catalytic PARP domain. Surprisingly, the structure revealed that the postulated binding site for the co-substrate NAD^+^ substantially differs from the structurally conserved NAD^+^ binding sites of canonical PARP enzymes. Consistently, we find that Ta-sro1 does not bind NAD^+^ and lacks ADP-ribosyltransferase activity. Therefore, although the structure revealed that one of the polymorphic amino acids is located close to the proposed active site, the elevated salinity tolerance of cultivar SR3 cannot be explained by altered ADP-ribosyltransferase activity of Ta-sro1.

## Introduction

High soil salinity affects 6% of the world’s land area and poses a significant limitation to agricultural production on approximately 20% of the irrigated farm land (Munns & Tester, 2008). Two components of high soil salinity affect plant growth. First, the lower water potential of saline soils aggravates water uptake by the root system causing loss of turgor pressure and closure of stomata. Secondly, long-term exposure to saline soils promotes accumulation of high Na^+^ concentrations in the shoot where Na^+^ ions interfere with photosynthesis leading to early senescence (Lawlor & Cornic, 2002; Munns & Tester, 2008; Chaves *et al*., 2009). Plant species substantially differ in their salt tolerance. Growth of glycophytes like rice and *Arabidopsis thaliana* (hereafter Arabidopsis) is strongly impaired by NaCl concentrations above 80 mM. Bread wheat and barley show intermediate salinity tolerance, whereas halophytes like tall wheat grass (*Thinopyrum ponticum*) continue to grow at NaCl concentrations equivalent to those in sea water (0.6 M) (Munns & Tester, 2008).

Plant strategies to cope with the ionic component of high salinity stress include active export of Na^+^ ions to the rhizosphere, control of Na^+^ loading and unloading of the xylem, and sequestration of Na^+^ in vacuoles (Julkowska & Testerink, 2015; Ismail & Horie, 2017). Plant cells sense high salinity by the osmotic changes and the increased concentration of Na^+^ in the extracellular space (Yuan *et al*., 2014; Julkowska & Testerink, 2015; Jiang *et al*., 2019; Lamers *et al*., 2020). Ca^2+^ acts as a second messenger in salt stress signaling and Ca^2+^ signatures are decoded by several calcineurin B-like proteins (CBLs) that cooperate with CBL-interacting protein kinases (CIPKs) to relay Ca^2+^ signals into phosphorylation of target proteins such as the plasma membrane localized Na^+^/H^+^ antiporter SALT OVERLY SENSITIVE 1 (SOS1) (Halfter *et al*., 2000; Shi *et al*., 2000; Choi *et al*., 2014; Manishankar *et al*., 2018). Salt stress also induces elevated production of reactive oxygen species (ROS) (Xie *et al*., 2011). ROS and Ca^2+^ signals can re-enforce each other and mathematical modeling suggests that interdependence of Ca^2+^ and ROS signaling underlie long-distance signaling in response to salt stress (Dubiella *et al*., 2013; Gilroy *et al*., 2014; Evans *et al*., 2016). One source of stress-induced ROS production in plants are proteins of the Respiratory Burst Oxidase Homolog (RBOH) family (Torres *et al*., 2002; Choudhury *et al*., 2017). An Arabidopsis *rbohD rbohF* double mutant is hypersensitive to high salinity stress and shows an attenuated NaCl-induced increase in cytosolic Ca^2+^ (Ma *et al*., 2012). In addition to their function as signal transducers, ROS can also become toxic to cells if they accumulate to high levels (Choudhury *et al*., 2017). Therefore, plants employ several ROS-scavenging system to prevent excessive oxidative damage of cellular components (Mittler *et al*., 2004). Collectively, orchestration of early signaling events by Ca^2+^ and ROS leads to establishment of mechanisms that counteract the osmotic and ionic components of high salinity stress (Julkowska & Testerink, 2015; Ismail & Horie, 2017).

Bread wheat cultivar Shanrong No. 3 (SR3), the product of an asymmetric genome hybridization between *Triticum aestivum* cultivar Jinan 177 (JN177) and the halophyte *T. ponticum*, exhibits elevated salinity stress tolerance compared to its parental line JN177. The superior salinity tolerance of cultivar SR3 has been linked to two amino acid polymorphisms in the *T. aestivum* SRO1 (SIMILAR TO RCD ONE 1) protein (Liu *et al*., 2014). SRO proteins with a WWE-PARP-RST (domain with conserved Trp/Glu residues / poly(ADP-ribose) polymerase(-like) / RCD1-SRO-TAF4) domain structure from different plant species have been genetically linked to altered abiotic stress tolerance (Ahlfors *et al*., 2004; You *et al*., 2013, 2014; Hiltscher *et al*., 2014). SRO proteins interact with transcription factors via their C-terminal RST domain and function as transcriptional co-regulators (Jaspers *et al*., 2009; Vainonen *et al*., 2016; Shapiguzov *et al*., 2019). The best-characterized SRO protein is RADICAL-INDUCED CELL DEATH 1 (RCD1) from *Arabidopsis thaliana*. Loss of *RCD1* function results in a pleiotropic phenotype and genes that are differentially regulated in *rcd1* mutants belong to several functional categories including redox homeostasis, innate immunity, programmed cell death, response to phytohormones, and development (Jaspers *et al*., 2009; Brosché *et al*., 2014; Wirthmueller *et al*., 2018). *rcd1* mutants are more tolerant to oxidative stress induced in chloroplasts (Ahlfors *et al*., 2004; Hiltscher *et al*., 2014) and a molecular link between RCD1 protein function and oxidative stress signaling has recently been established. RCD1 interacts with two NAC transcription factors that promote expression of mitochondrial dysfunction stimulon (MDS) genes in response to mitochondrial ROS (Shapiguzov *et al*., 2019). In *rcd1* mutants, MDS genes including alternative oxidase genes are constitutively expressed and this mitigates toxic effects of ROS produced in chloroplasts (Shapiguzov *et al*., 2020). Retrograde signaling from mitochondria and chloroplasts to the nucleus is altered in *rcd1* mutants and ROS produced in chloroplasts affect abundance, oligomerization and redox state of the nuclear RCD1 protein (Shapiguzov *et al*., 2019). Results from Arabidopsis and other plant species suggest that an altered redox homeostasis is a common consequence of mutations in *SRO* genes and this could affect plant responses to different types of biotic and abiotic stresses such as high salinity (You *et al*., 2013, 2014; Liu *et al*., 2014; Brosché *et al*., 2014; Wirthmueller *et al*., 2018; Shapiguzov *et al*., 2019; Zhang *et al*., 2019).

The RST domain of SRO proteins forms a four helix bundle that directly interacts with transcription factors (Jaspers *et al*., 2009, 2010; Kjaersgaard *et al*., 2011; O’Shea *et al*., 2015; Bugge *et al*., 2018; Shapiguzov *et al*., 2019). In contrast, the roles of the WWE and PARP domains remain largely unknown. Recent results suggest that, similar to several mammalian WWE domains, the WWE domain of RCD1 binds to poly(ADP-ribose) (PAR) chains *in vitro* (Vainonen *et al*., 2021). The RCD1 WWE and PARP domains are both required for localization of RCD1 in nuclear bodies suggesting that these domains control the subnuclear location of RCD1, possibly in dependence on interaction with PAR chains (Vainonen *et al*., 2021). The central PARP domain was named based on sequence homology to the catalytic domains of canonical PARPs, enzymes that use NAD^+^ as a co-substrate to transfer ADP-ribose moieties onto themselves and other target proteins (Vainonen *et al*., 2016; Rissel & Peiter, 2019). The ADP-ribosyltransferase activity of canonical PARPs is not limited to amino acids. Following the initial modification of an amino acid side chain, certain PARPs can attach further ADP-ribose moieties onto the terminal ADP-ribose thereby forming PAR chains (Hottiger, 2015). A structure of the Arabidopsis RCD1 PARP domain revealed that three amino acids that mediate NAD^+^ binding and catalysis in canonical PARPs are not conserved in RCD1 (Wirthmueller *et al*., 2018). Accordingly, RCD1 does not bind NAD^+^ and mutations at the presumed active site do not affect RCD1 function in Arabidopsis (Jaspers *et al*., 2010; Wirthmueller *et al*., 2018). Based on sequence alignments and homology modeling, the catalytic His-Tyr-Glu triad of canonical PARPs is not conserved in SRO proteins and this includes wheat SRO1 (Ta-SRO1) (Jaspers *et al*., 2010; Liu *et al*., 2014; Vainonen *et al*., 2016). Surprisingly, the Ta-SRO1 protein retains PARP activity despite non-conservation of the catalytic triad (Liu *et al*., 2014). Moreover, Liu *et al*. (2014) proposed that the superior salinity stress tolerance of wheat cultivar SR3 is largely explained by the elevated PARP activity of the hypermorphic Ta-sro1 proteoform. To understand why Ta-sro1 retains PARP activity despite the lack of the catalytic triad we solved a crystal structure of the PARP domain and biochemically characterized the postulated Ta-sro1 PARP activity.

## Materials and Methods

### Molecular cloning

To generate His_6_-tagged expression constructs, the coding sequences of Ta-sro1 WWE-PARP (residues 2-434) and Ta-sro1 PARP (residues 246-434) were cloned into *Kpn*I/*Hind*III-linearized pOPIN-F (Berrow *et al*., 2007) via Gibson assembly. The pET24 plasmid for expression of full-length Ta-sro1 was a kind gift from Prof. Guangmin Xia (Liu *et al*., 2014). The expression constructs for the catalytic domains of HsPARP1 and HsPARP10 have been described (Kleine *et al*., 2008; Langelier *et al*., 2012). For transient expression in *Nicotiana benthamiana*, the following coding sequences were cloned into *Nco*I/*Xho*I-linearized pENTR4 via Gibson assembly. Ta-sro1 and Ta-SRO1 (residues 1-578, no stop codon), Ta-sro1 WWE-PARP (residues 1-434, no stop codon), human RNF146 WWE domain (residues 99-183, including stop codon). The pENTR4 AtPARP2 plasmid has been described (Chen *et al*., 2018). The AtPARP2 point mutation E^614^Q and the RNF146 Tyr^156^/Arg^157^ to Ala mutations were introduced in pENTR4 by site-directed mutagenesis. pENTR4-GFP-C3 (w393-1) was a gift from Eric Campeau & Paul Kaufman (Addgene plasmid # 17397; http://n2t.net/addgene:17397; RRID:Addgene_17397). The pENTR4 plasmids were recombined in Gateway LR reactions with pGWB414 (Nakagawa *et al*., 2007) to create expression constructs with a C-terminal 3xHA tag. To create expression constructs with a C-terminal eGFP tag, the respective pENTR4 plasmids were recombined in Gateway LR reactions with pK7FWG2 (Karimi *et al*., 2002). For RFP-tagged constructs the respective pENTR4 plasmids were recombined in Gateway LR reactions with pH7WGR2 (N-terminal RFP tag) or pH7RWG2 (C-terminal RFP tag) (Karimi *et al*., 2002). Oligonucleotide sequences are listed in Table S2.

### Protein expression and purification

For small-scale protein expression trials *in Escherichia coli*, overnight cultures of the respective expression clones were diluted 1:60 in 3 mL LB medium with appropriate antibiotics in 24-well plates. The 3 mL cultures were incubated on a platform shaker at 37 °C and 130 rpm until the OD_600_ reached 0.8 – 1. The temperature was lowered to 18 °C before protein expression was induced by addition of IPTG to a final concentration of 0.5 mM. After 16 h, the 3 mL cultures were pelleted and the cells were resuspended in 1 mL lysis buffer [50 mM Tris-HCl pH 8.0, 0.3 M NaCl, 50 mM glycine, 5% glycerol, 0.1% Polyethylenimine (Merck), 0.1% Triton X-100, 0.05 mg/mL DNAse I (Applichem), 0.5 mg/mL Lysozyme, 1x cOmplete™ EDTA-free protease inhibitor cocktail (Merck)]. Following incubation on a platform shaker (200 rpm / 25 °C / 10 min), 0.2 mL of the cell suspension was transferred to a 1.5 mL tube and sonicated for 2x 10 min in a Bandelin Sonorex Digitec DT 514 H sonication bath. A fraction of the lysate was saved as ’total’ protein sample and the remaining lysate was centrifuged at 20.000 x *g* / 25 °C / 5 min and the supernatant saved as ’soluble’ protein sample.

The Ta-sro1-His_6_ full-length protein was expressed from pET24a in *E. coli* BL21 cells (Liu *et al*., 2014). The Ta-sro1 His_6_-WWE-PARP and His_6_-PARP constructs were expressed from pOPIN-F in *E. coli* SHuffle cells. The HsPARP1 catalytic domain construct was expressed from pET28 in *E. coli* SoluBL21 cells (Langelier *et al*., 2012). The GST-HsPARP10 catalytic domain containing amino acids 818-1025 was produced in *E. coli* BL21 cells essentially as described before (Kleine *et al*., 2008), with induction of protein expression at an OD_600_ of 0.6 with 1 mM IPTG, followed by 16 h incubation at 18 °C. Bacterial cultures were spun down (6000 x *g* / 4 °C / 15 min), followed by resuspension in lysis buffer [20 mM Tris pH 8.0, 150 mM NaCl, 0.1 mM EDTA pH 8.0, 5 mM DTT and 1x protease inhibitor cocktail (Sigma)], supplemented with 1 mg/mL Lysozyme (final concentration) and incubated on ice for 30 min. Cell suspensions were sonicated on ice (Branson Digital Sonifier 250 Cell Disruptor, 5 min at 20%, 30 s on/off rate). The cell lysates were centrifuged at 45,000 x *g* for 45 min at 4°C to remove cell debris. The supernatant was used for affinity purification and incubated with Glutathione Sepharose 4B (GE healthcare). The beads were washed in wash buffer (100 mM Tris pH 8.0, 120 mM NaCl) followed by elution using glutathione (20 mM in wash buffer, prepared fresh) and dialysis overnight [20 mM Tris pH 8.0, 150 mM NaCl, 1 mM DTT, 10% (v/v) glycerol].

For Ta-sro1 expression constructs, two to eight 1 L cultures were grown in LB medium at a temperature of 37 °C to an OD_600_ of 1.0 – 1.2. The cultures were cooled to 18 °C before expression was induced by the addition of 0.5 mM IPTG for 16 h. Cells were pelleted by centrifugation (5000 x *g* / 4 °C / 12 min) and the pellets were resuspended in buffer A [50 mM Tris-HCl pH 8.0, 0.3 M NaCl, 20 mM imidazole, 5% (v/v) glycerol, 50 mM glycine] supplemented with 0.1% Polyethylenimine and 1x cOmplete™ EDTA-free protease inhibitor cocktail (Merck). Cells lysis was induced by addition of Lysozyme (1 mg/mL final concentration / 25 °C / 15 min) followed by sonication on ice (Branson 150D Sonifier, 2x 10 min, level 3-4). Insoluble proteins and cell debris were removed by centrifugation (30.000 x *g* / 4 °C / 30 min) and the supernatant was loaded onto a 5 mL HisTrap HP IMAC column (Cytiva). The column was washed with buffer A until the A_280_ reached 25 mAU and proteins were eluted using buffer B [50 mM Tris-HCl pH 8.0, 0.3 M NaCl, 500 mM imidazole, 5% (v/v) glycerol, 50 mM glycine]. The elution from the IMAC column was injected onto a size exclusion chromatography column [Superdex 75 or Superdex 200 26/60 PG column (Cytiva) pre-equilibrated with 20 mM HEPES-NaOH pH 7.5, 150 mM NaCl]. Proteins eluting from the column were concentrated by ultrafiltration on Vivaspin 20 and 2 columns (Sartorius) with a 5 kDa molecular weight cut-off. The Selenomethionine-labeled Ta-sro1 PARP domain was produced using feedback inhibition (Van Duyne *et al*., 1993) and purified as described above. For crystallization, the His_6_-tag was cleaved using 3C protease. The protein was run through a 5 mL HisTrap HP IMAC column in buffer A to remove the His_6_-tag and residual un-cleaved fusion protein, followed by injection onto the Superdex 75 26/60 PG column and eluted and concentrated as above.

### Protein crystallization and structure determination

Crystals of native and Selenomethionine-labeled PARP domain formed in 0.1 M MES-KOH pH 5.8, 0.2 M ammonium sulfate, 17% (w/v) PEG 3350 at a protein concentration of 23.3 mg/mL in a vapor diffusion setup at 291 K. The crystals were transferred to cryo protectant solution [0.1 M MES-KOH pH 5.8, 0.2 M ammonium sulfate, 17% (w/v) PEG 3350, 25% (v/v) ethylene glycol] and flash-frozen in liquid nitrogen. Diffraction data was collected at beamline 14.2 of the BESSY II synchrotron, Berlin at 100 K. The data were processed with XDS (Kabsch, 2010; Sparta *et al*., 2016) and the PARP domain structure was solved by single anomalous diffraction phasing using the AutoSol wizard in PHENIX (Adams *et al*., 2010). The final model was obtained by iterative building and refinement cycles using Coot (Emsley *et al*., 2010) and PHENIX (Afonine *et al*., 2012). In final stages of refinement, TLS refinement (Winn *et al*., 2001) was performed as implemented in PHENIX. The model was validated using Molprobity (Chen *et al*., 2010) and Coot (Emsley *et al*., 2010). The statistics for X-ray data collection and the refined model are given in Table S1. Reflection data and the Ta-sro1 PARP domain structure have been deposited at the Protein Data Bank with identifier 7PLQ. Diffraction images have been deposited at www.proteindiffraction.org under DOI:10.18430/M37PLO. 3D visualizations of protein structures were prepared using PyMol software v1.7.2 (https://sourceforge.net/projects/pymol/).

### Analytical size exclusion chromatography

50 μg of Ta-sro1 WWE-PARP protein was diluted in 700 μL of SEC buffer (20 mM HEPES-NaOH pH 7.5, 150 mM NaCl) and injected onto a Superdex 200 Increase 10/300 GL column using a 380 μL sample loop. For other experiments, the protein was pre-incubated in SEC buffer supplemented with NaCl (final concentration 0.5 M) or DTT (final concentration 10 mM) for 30 min at RT. Before each run, the SEC column was equilibrated with two column volumes of the respective buffer. To test for PAR-binding, 25 μg of Ta-sro1 WWE-PARP was mixed with 175 μL SEC buffer and 150 μL 10 μM PAR (Trevigen). The control sample consisted of 25 μg of Ta-sro1 WWE-PARP and 325 μL SEC buffer.

### Thermal stability assay

Thermal stability assays were performed in a MX3005 qPCR System (Agilent Technologies) with thermal denaturation ranging from 25 °C to 95 °C in 0.5 °C steps. All samples were measured as technical triplicates and contained 25 mM NaCl, 75 mM Tris-HCl pH 7.5 and 6.6x SYPRO™ Orange (Invitrogen) in a final volume of 20 µL. Negative controls additionally contained the highest concentration of ligand used [2 mM 6(5H)-phenanthridinone or 4 mM NAD^+^], but no protein or only protein without ligand. All other samples contained the same amount of protein (0.11 mg/mL) and 7 ligand concentrations ranging from 4 nM to 4 mM for NAD^+^ and 2 nM to 2 mM for 6(5H)-phenanthridinone. The global minimum of the negative derivative of fluorescence over temperature was considered the melting temperature. Melting temperatures were averaged from three technical replicates and three independent experiments and were plotted against ligand concentrations.

### NAD^+^ binding and auto-ADP-ribosylation assays

NAD^+^ binding was tested by spotting the proteins on nitrocellulose membrane followed by incubation with radiolabelled NAD^+^. Serial dilutions of the proteins were spotted, with 500 ng protein as highest concentration followed by 1:2 dilutions. After spotting the proteins, the membrane was allowed to dry before blocking in 5% non-fat milk in PBST. The blocked membrane was washed extensively in PBST before addition of 10 μCi [^32^P]-β-NAD^+^ (Hartmann Analytic) in PBST. After one h incubation at RT, the membrane was washed again in PBST, followed by exposure to X-ray film of the dried membrane. HsPARP10cat served as positive control, GST as negative control. Protein auto-ADP-ribosylation assays were performed at 37 °C for 30 min. Reactions were carried out in 30 µL volume containing 50 mM Tris-HCl pH 8.0, 0.2 mM DTT, 4 mM MgCl_2_ and 50 μM β-NAD^+^ (Sigma) and 1 μCi [^32^P]-β-NAD^+^ (Hartmann Analytic). Reactions were stopped by adding SDS sample buffer, heated for 5 min at 95 °C and analyzed using SDS-PAGE. Gels were dried and incorporated radioactivity was analyzed by exposure of the dried gel to X-ray film. HsPARP10cat served as positive control, GST as negative control.

### PARylation assay

The PARP activity assay was performed with the PARP Universal Colorimetric Assay Kit (Trevigen) according to the manufacturer’s manual, section ’PARP Inhibitor Assay Protocol’. 0.5 μg of protein was added per well and each reaction was performed in triplicate. Within the recommended assay time of 1 h, the two HsPARP1 reactions produced A_450_ values that were too high to be measured by the Tecan Infinite F50 plate reader. Therefore, the reactions were already stopped and quantified after 15 min.

### *Agrobacterium*-mediated transient expression

Binary vectors were transformed into *Agrobacterium tumefaciens* strain GV3101 pMP90. *A. tumefaciens* strains were grown on selective LB plates, resuspended in 10 mM MgCl_2_ 10 mM MES-KOH pH 5.6 and incubated with 100 μM acetosyringone for 2 h at RT. Prior to infiltration, each strain was mixed with *A. tumefaciens* strain GV3101 pMP90 expressing the silencing suppressor 19K at a ratio of 1:1.5[19K]. For co-expression, strains were mixed in a 1:1:1.5[19K] ratio. The cultures were infiltrated into leaves of 4-5 week-old *N. benthamiana* plants using a needleless syringe and leaf material for protein extraction or confocal microscopy was harvested 48–72 h later.

### Confocal microscopy

Leaf discs excised from *N. benthamiana* were mounted on microscope slides in 50% (v/v) glycerol and the subcellular localization of GFP-tagged proteins was analyzed using a Zeiss LSM 780 confocal microscope. Excitation wavelength and emission collection windows were 488/493-545 nm for GFP and 561/687-730 nm for chlorophyll. Images were acquired and analyzed using Zeiss Zen software.

### Protein extraction and immunopurification from leaf tissue

Protein extracts were prepared by grinding *N. benthamiana* leaf material in liquid nitrogen to a fine powder followed by resuspension in extraction buffer [50 mM Tris-HCL pH7.4, 150 mM NaCl, 10% (v/v) glycerol, 1 mM EDTA, 5 mM DTT, 1× protease inhibitor cocktail (Merck #P9599), 0.2% NP-40] at a ratio of 2 mL buffer per 1 g leaf material. In assays to detect protein (auto-)ADP-ribosylation, the extraction buffer was supplemented with 300 μM NAD^+^. Crude protein extracts were centrifuged at 20.000 x *g* / 4 °C / 20 min and the supernatant was either boiled in sodium dodecyl sulphate (SDS) sample buffer for western blots or used for immunoprecipitation. For immunoprecipitation a fraction of the supernatant was saved as ‘input’ sample and 15 μL of α-GFP-nanobody:Halo:His_6_ magnetic beads (Chen *et al*., 2018) were added to 1.4 mL of the remaining supernatant. The samples were incubated on a rotating wheel at 4 °C for 2 h followed by collection of the beads using a magnetic sample tube rack. The beads were washed 3 times with 1 mL extraction buffer and then boiled in 40 μL SDS sample buffer to elute protein from the beads.

### SDS-PAGE and immunoblotting

Protein samples were separated by SDS-PAGE and stained using Instant Blue (Thermo-Fisher). For SDS-PAGE under non-reducing conditions, the proteins were boiled in SDS sample buffer without DTT. For immunoblots, proteins were electro-blotted onto PVDF membrane and blocked with a solution of 5% non-fat dry milk in TBST for 1 h at RT. Antibodies used were α-HA 3F10 (Merck), α- GFP TP401 (Amsbio), Biotin α-RFP ab34771 (Abcam), α-His_6_ (anti-polyhistidine tag antibody, Merck), and α-MAR/PAR (anti-pan-ADP-ribose binding reagent, Merck). Proteins were detected using HRP-coupled secondary antibodies and X-ray films.

### Sequence analysis and alignments

Disordered regions of Ta-sro1 were predicted using the PONDR VL3-BA algorithm (Xue *et al*., 2010). DNA and protein sequences were aligned with Clustal Omega (Sievers *et al*., 2011).

## Results

### The presumed active site of Ta-sro1 differs in several key features from canonical PARPs

To understand the structural basis for the non-canonical PARP activity of Ta-sro1 we expressed His_6_-tagged variants of the full-length protein as well as the PARP domain in *E. coli* (Fig. 1a; Fig. S1a, b). The purified full-length Ta-sro1 protein eluted in two peaks from a size exclusion chromatography (SEC) column and showed signs of protein degradation (Fig. S1c). In contrast, the isolated PARP domain was stable and crystallized readily. We solved the structure of the Selenomethionine-labeled PARP domain by single anomalous diffraction phasing at 2.1 Å resolution (Table S1, Fig. S1d). The domain adopts the typical PARP fold with an β-α−loop-β-α signature at the donor site that binds NAD^+^ (Fig. 1b). The donor site loop (D-loop) occludes the proposed active site of Ta-sro1 and is only partially resolved in the structure with little interpretable electron density for amino acids Met^334^ – Gly^336^ (Fig. 1b). A structural homology search using the Dali server (Holm, 2020) identified PARP domains of the Arabidopsis paralog RCD1 and human PARP10 and PARP13 proteins as closest homologs (Z scores 18.2 – 22.1). HsPARP10 is limited to mono-ADP-ribosyl transferase activity whereas HsPARP13 has lost the ability to bind the co-substrate NAD^+^ (Kleine *et al*., 2008; Karlberg *et al*., 2015). The structure confirms that three key catalytic residues of canonical PARP enzymes are not conserved in Ta-sro1 (Fig. 1b). The position of the conserved His residue that forms hydrogen bonds with the proximal NAD^+^ ribose is taken by Ta-sro1 Leu^312^. The Tyr that stacks with the distal ribose ring in canonical PARP enzymes is replaced by His^344^. The catalytic Glu that is specifically required for the PAR chain elongating polymerase activity of PARPs is replaced by Ta-sro1 His^407^. The Ta-sro1 PARP domain structure pinpoints the location of the two amino acid polymorphisms that distinguish the hypermorphic proteoform Ta-sro1 from Ta-SRO1 and are responsible for the enhanced abiotic stress tolerance (Liu *et al*., 2014). Ta-sro1 Thr^343^ (Ala in Ta-SRO1) is located close to the proposed active site and its side chain is positioned towards the hydrophobic core of the protein formed by residues Leu^345^, Val^330^, Leu^328^, Ile^419^, Val^415^, and Ile^410^ (Fig. 1b, c). The second polymorphic amino acid, Val^250^ (Gly in Ta-SRO1) is positioned on the opposite side of the domain and located close to the N-terminus of the crystallized construct (Fig. 1b). In context of the full-length protein, Val^250^ would form part of the transition from the predicted intrinsically disordered region 2 (IDR2) to the PARP domain (Fig. 1a). Understanding the functional relevance of the Gly^250^ to Val exchange therefore may require further structural information in context of the WWE-PARP domains. Despite the altered identity of amino acids that form the NAD^+^-binding site, Ta-sro1 apparently can still utilize NAD^+^ as a co-substrate (Liu *et al*., 2014). To understand how Ta-sro1 makes contact to NAD^+^ we superimposed the Ta-sro1 PARP domain onto the structure of the catalytic domain of human PARP1 crystallized in complex with the NAD^+^ analog benzamide adenine dinucleotide (PDB identifier 6BHV) (Langelier *et al*., 2018). The two structures superimposed with a root-mean-square deviation of 2.28 Å over 135 aligned residues. Fig. 1d shows the active site of HsPARP1 (grey) in comparison to the presumed Ta-sro1 active site (beige). In addition to the altered identity of several residues that make contact with NAD^+^ (described above), it is apparent that the side chain of Pro^313^ would sterically interfere with NAD^+^ binding if Ta-sro1 would bind the co-substrate in the same orientation as canonical PARPs. The corresponding residue in catalytically active mono- and poly-ADP-ribosyltransferases is a conserved Gly that accommodates the amide group of the nicotinamide moiety and stabilizes it by forming two hydrogen bonds (Wahlberg *et al*., 2012). In contrast, a second Tyr that is positioned on the opposite side of the cleft, stacking with the nicotinamide ring, is conserved in Ta-sro1 (Tyr^357^) (Fig. 1d). Overall, the PARP domain structure reveals that the presumed active site of Ta-sro1 substantially differs in several key residues from canonical PARP domains of plants and mammals (Wahlberg *et al*., 2012; Vyas *et al*., 2014; Vainonen *et al*., 2016; Gu *et al*., 2019).

**Figure 1.**
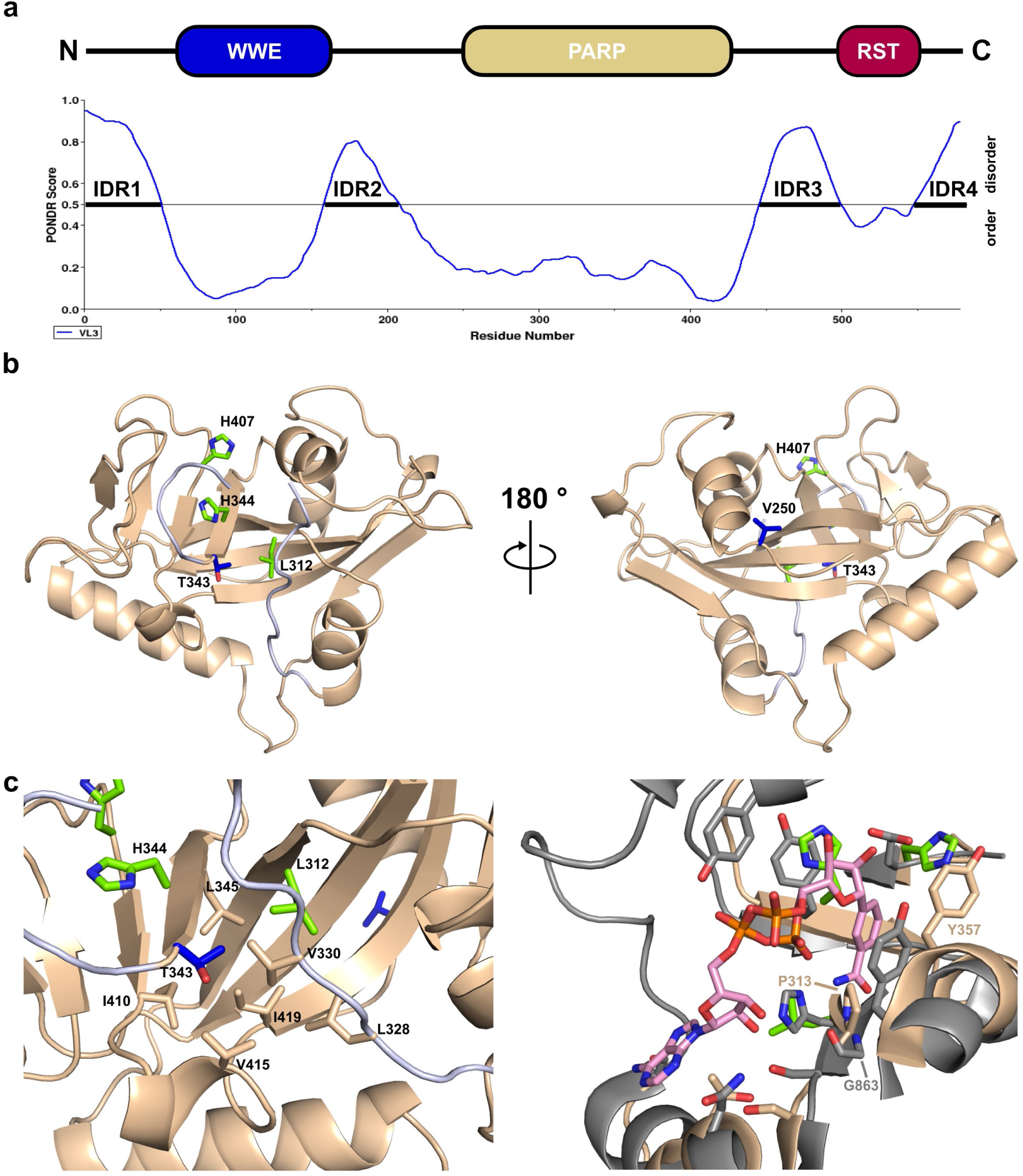
Crystal structure of the Ta-sro1 PARP domain. **(a)** Domain architecture and predicted protein disorder profile of Ta-sro1. Intrinsically disordered regions are labelled IDR1-4. **(b)** Structure of the Ta-sro1 PARP domain (residues 247 – 429) in cartoon representation. The postulated catalytic Leu-His-His triad is shown in green. Blue color indicates the two polymorphic residues Val^250^ and Thr^343^ that distinguish the two proteoforms. The partially disordered D-loop is shown in grey. **(c)** The side chain of the polymorphic residue Thr^343^ does not contribute to the postulated NAD^+^ binding site but points towards the hydrophobic core of the PARP domain. **(d)** Superposition of the Ta-sro1 PARP domain (beige) and the HsPARP1 PARP domain (grey) co-crystallized in complex with the NAD^+^ analog benzamide adenine dinucleotide (PDB identifier 6BHV). The position taken by Ta-sro1 residue Pro^313^ is an invariant Gly in catalytically active PARPs and Pro^313^ would sterically interfere with binding of the co-substrate NAD^+^.

### Ta-sro1 is catalytically inactive with respect to canonical ADP-ribosylation

The observed structural divergence from the consensus motif at the active site of the Ta-sro1 PARP domain prompted us to test whether Ta-sro1 can indeed bind NAD^+^ and is able to perform ADP-ribosylation reactions. For mammalian PARP domains, thermal stabilization of the catalytic domain by small molecule inhibitors that mimic nicotinamide can serve as a proxy for NAD^+^ binding (Wahlberg *et al*., 2012). We determined the thermal stability of the Ta-sro1 PARP domain with increasing concentrations of the nicotinamide analog 6(5H)-phenanthridinone that was previously shown to stabilize ten catalytically active mammalian PARP domains (Wahlberg *et al*., 2012). We did not observe a thermal stabilization of the Ta-sro1 PARP domain by 6(5H)- phenanthridinone, even at concentrations in the millimolar range (Fig. 2a). In contrast, a variant of the HsPARP1 catalytic domain (Langelier *et al*., 2012) showed increased thermal stability (∼+4 °C) at 6(5H)-phenanthridinone concentrations above 2 μM (Fig. 2a). Morevover, in thermal stability assays with NAD^+^ as a ligand, we did not observe an altered unfolding temperature of the Ta-sro1 PAPR domain. For the HsPARP1 catalytic domain there was only a marginal stabilization at NAD^+^ concentrations above 40 μM (Fig. 2a) but the absence of a stabilizing effect at lower concentrations is likely due to consumption of NAD^+^ by auto-ADP-ribosylation during preparation of the reaction mixture.

**Figure 2.**
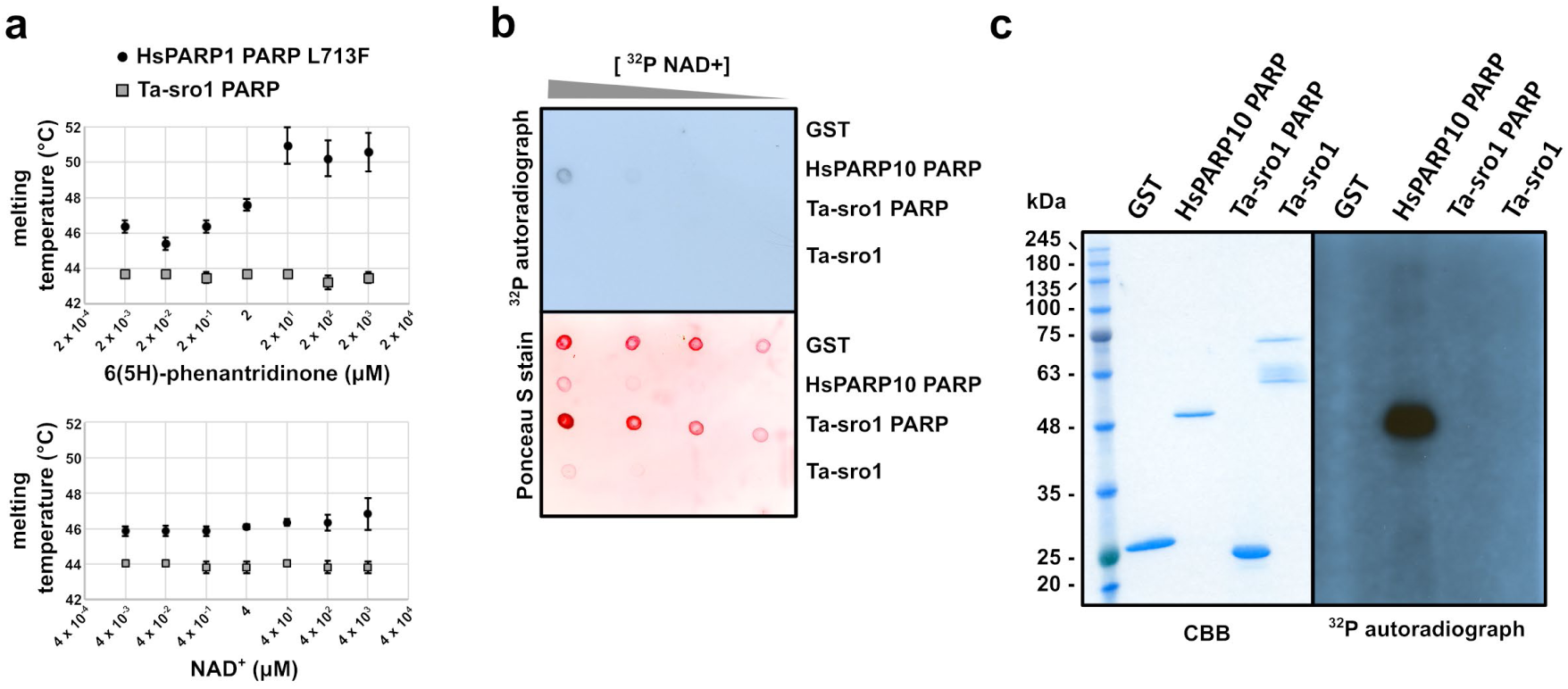
The Ta-sro1 PARP domain does not bind NAD^+^ and does not auto-ADP-ribosylate *in vitro*. **(a)** Thermal stability profiles of the Ta-sro1 PARP domain and the hypermorphic L^713^F variant of the HsPARP1 PARP domain. Calculated domain melting temperatures were averaged from three independent experiments, each consisting of three technical replicates per sample. Melting temperatures are plotted over a concentration range of the NAD^+^ analog 6(5H)-phenantridinone (top) or NAD^+^ (bottom). Error bars represent standard deviations. **(b)** NAD^+^ binding was tested by spotting serial dilutions of the indicated proteins on nitrocellulose membrane followed by incubation with radiolabelled NAD^+^ and exposure to X-ray film. The HsPARP10 PARP domain served as positive control, GST as negative control. The Ponceau S stain indicates protein amounts spotted onto the membrane. **(c)** Protein auto-ADP-ribosylation assays were performed with indicated proteins at 30°C for 30 min and analyzed using SDS-PAGE. Gels were dried and incorporated radioactivity was analyzed by exposure of the dried gel to X-ray film. The HsPARP10 PARP domain served as positive control, GST as negative control. The results from (b) and (c) are representative of two independent experiments. CBB, Coomassie Brilliant Blue stain.

An *in vitro* binding assay with ^32^P-labeled NAD^+^ revealed that neither full-length Ta-sro1 nor the isolated PARP domain can bind NAD^+^ (Fig. 2b). In contrast, the same assay detected ^32^P-NAD^+^ binding to HsPARP10 that was used as a positive control. Many ADP-ribosyltransferases of the PARP family auto-ADP-ribosylate in the presence of NAD^+^ (Vyas *et al*., 2014). Unlike the catalytic domain of HsPARP10 that attaches a single APD-ribose onto Glu residues in an intermolecular reaction (Kleine *et al*., 2008), neither the Ta-sro1 PARP domain nor the full-length protein showed auto-ADP-ribosylation *in vitro* when incubated with ^32^P-labeled NAD^+^ (Fig. 2c). The postulated PARP activity of Ta-sro1 is based on a commercially available colorimetric assay that uses histones as substrates for ADP-ribosyltransferases (Liu *et al*., 2014). We repeated this assay (Fig. 3a) for full-length Ta-sro1, the isolated PARP domain, and a protein fragment that also includes the N-terminal WWE domain of the protein (Ta-sro1 WWE-PARP). Compared to the positive control included in the assay kit (full-length HsPARP1) and the variant HsPARP1 L713F catalytic domain produced in our laboratory, all three Ta-sro1 constructs were inactive with respect to PARP activity (Fig. 3a). Values for relative PARP activities of the negative control BSA and a reaction without protein were in the same range as those of the Ta-sro1 constructs. PARP activity of HsPARP1 was inhibited by the nicotinamide analog 3-aminobenzamide (3AB), demonstrating that the assay truly reflects PARP activity (Fig. 3a). Therefore, the standardized assay on which the previously reported Ta-sro1 PARP activity is based, is not reproducible under our conditions.

**Figure 3.**
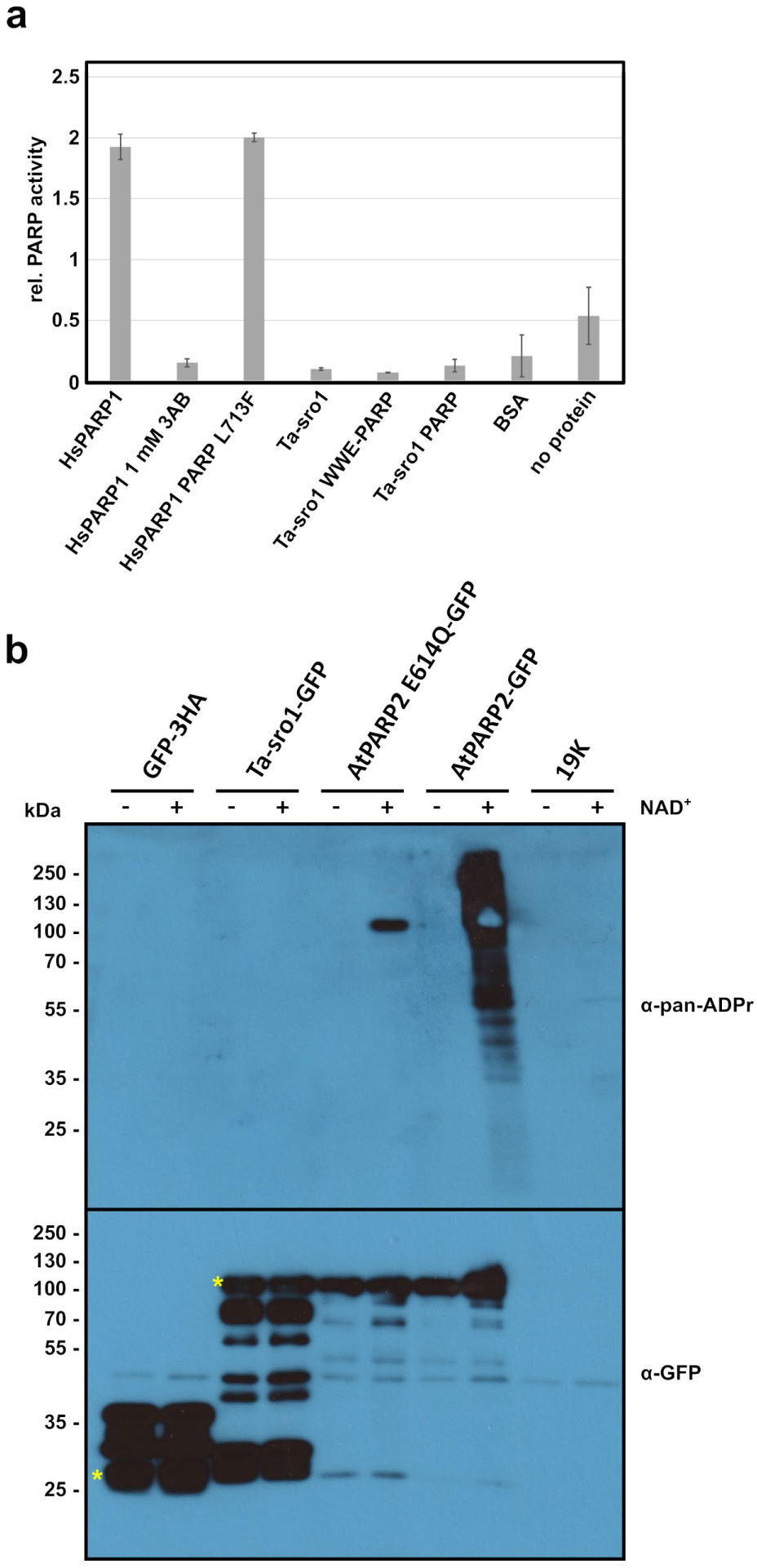
Ta-sro1 protein is catalytically inactive with respect to canonical mono-or poly-ADP- ribosylation. **(a)** Representative result of the colorimetric PARP activity assay using histones as general ADP-ribosylation substrates. HsPARP1 refers to activated full-length human PARP1 protein that is included as a positive control in the assay kit. Error bars represent standard deviations. **(b)** The indicated proteins were transiently expressed in *N. benthamiana* and extracted in presence (+) or absence (-) of 300 μM NAD^+^. Proteins were purified via the GFP-tag and the precipitated proteins were analyzed by α-GFP and α-pan-ADP-ribose binding reagent immunoblots. Leaves expressing only the viral silencing suppressor 19K or GFP-3HA served as negative controls. AtPARP2 and the AtPARP2-E^614^Q mutant represent controls for proteins with PARP and mono-ADP-ribosyltransferase activity, respectively. Asterisks indicate the expected molecular weights of GFP-3HA and GFP-fusions of Ta-sro1 and AtPARP2. The results from (a) and **(b)** are representative of three independent experiments.

Several PARP enzymes require other proteins for catalytic activity or depend on a binding partner to modify specific amino acids (Yang *et al*., 2017; Palazzo *et al*., 2018; Suskiewicz *et al*., 2020; Bilokapic *et al*., 2020). Given that the previously reported PARP activity of Ta-sro1 is based on an *in vitro* assay with recombinantly expressed protein, dependence on another plant factor to gain catalytic activity appears unlikely. Nevertheless, we assessed whether Ta-sro1 has mono- or poly-ADP-ribosyltransferase activity in plant cell extracts. We used transient Agrobacterium-mediated expression to produce Ta-sro1 and several control proteins as GFP fusions in *N. benthamiana*. We extracted the proteins in the absence or presence of 300 μM NAD^+^ and immunoprecipitated them using a GFP-nanobody coupled to magnetic beads (Kubala *et al*., 2010; Chen *et al*., 2018). After separation by SDS-PAGE, we probed soluble protein extracts with anti-pan-ADP-ribose binding reagent that binds to mono- and poly-(ADP-ribose) (Fig. 3b). Ta-sro1-GFP did neither show signals for mono- nor for poly-ADP-ribosylation activity in this assay. In contrast, the canonical PARP2 enzyme from Arabidopsis formed PAR chains in a NAD^+^-dependent manner. A PARP2 variant, in which the Glu residue that is essential for chain elongation is replaced by a Gln, produced a defined band of ∼100 kDa that could correspond to mono-ADP-ribosylated PARP2-GFP. These results suggest that even at relatively high concentrations of NAD^+^ and in presence of other plant co-factors that might be required for catalytic activity, Ta-sro1 does not exhibit mono- or poly-ADP-ribosylation activity.

### Ta-sro1 shows no detectable binding to PAR chains in plant cells

As shown by Vainonen *et al*. (2021), the Arabidopsis RCD1 WWE domain binds to PAR chains *in vitro* with a dissociation constant of ∼30 nM. To test whether also Ta-sro1 can bind to PAR chains and to probe for this interaction in plant cells, we established an *in planta* PAR chain binding assay. We used transiently expressed auto-PARylated Arabidopsis PARP2-GFP (Fig. 3b) as a bait to co-immunoprecipitate PAR-binding proteins from *N. benthamiana* leaf extracts. As a positive control for a PAR-binding protein, we used the human RNF146 WWE domain (Zhang *et al*., 2011; Andrabi *et al*., 2011; Wang *et al*., 2012). As shown in Fig. 4, PARylated PARP2-GFP co-immunoprecipitated the positive control RFP-WWE^RNF146^ but not free RFP or a mutated RFP-WWE^RNF146^ protein (Tyr^156^/Arg^157^ to Ala, Andrabi *et al*., 2011). In contrast, we could not detect strong interactions between auto-PARylated PARP2-GFP and the Ta-sro1 WWE-PARP fragment or the Ta-sro1 full-length protein fused to RFP. Therefore, either the ability to bind to PAR is not conserved in Ta-sro1 or the interaction is substantially weaker than the PAR/WWE^RNF146^ interaction.

**Figure 4.**
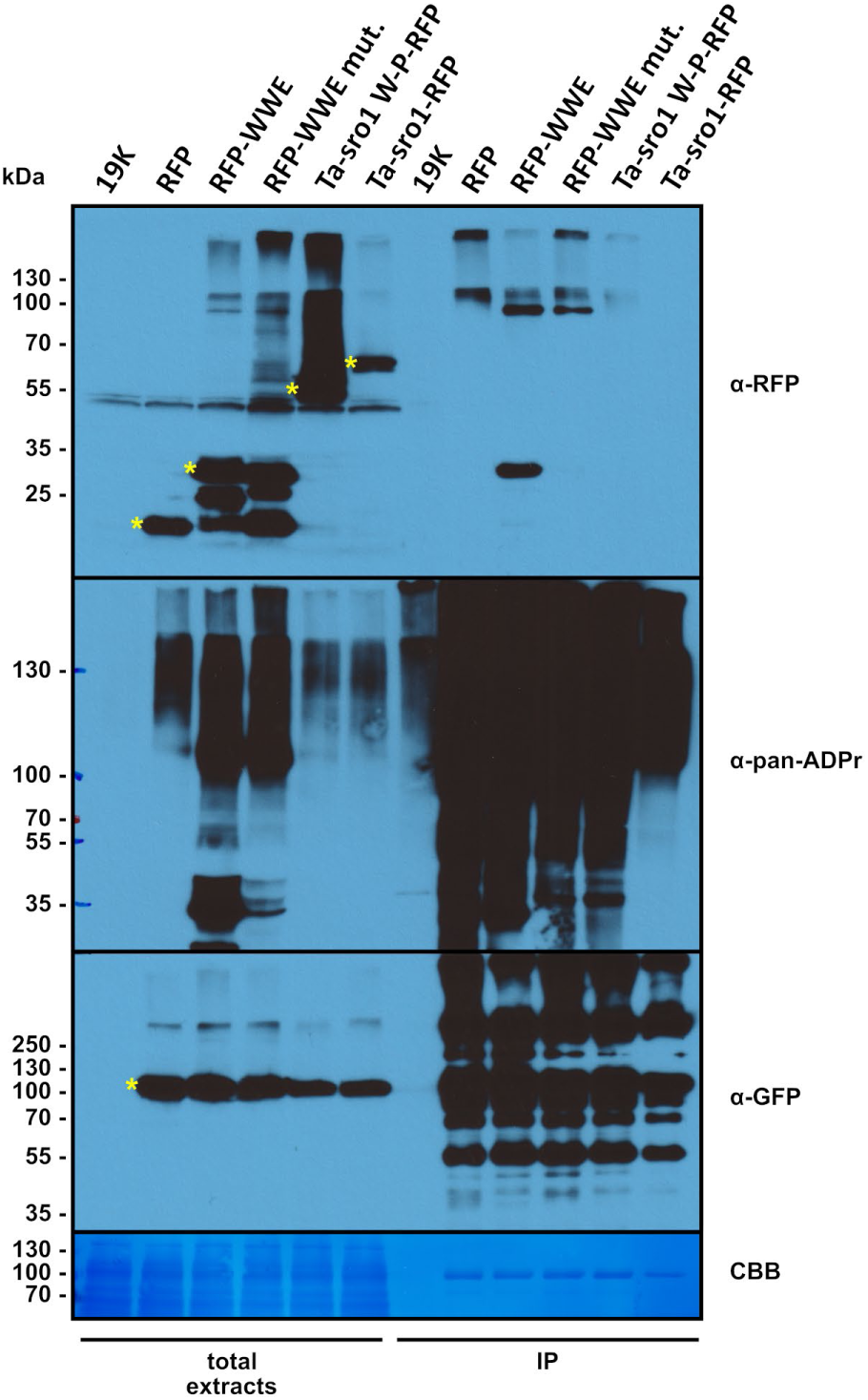
The Ta-sro1 WWE domain does not show strong interaction with PAR chains in plant cells. The indicated RFP-tagged proteins were co-expressed with AtPARP2-GFP in *N. benthamiana* and extracted in presence of 300 μM NAD^+^ to allow for AtPARP2 auto-PARylation. Leaves expressing only the viral silencing suppressor 19K served as control for endogenous PARylation from *N. benthamiana*. PARylated AtPARP2-GFP was purified via the GFP-tag and co-purifying RFP-tagged proteins were detected by an α-RFP immunoblot. RFP-WWE refers to the human RNF146 WWE domain that has been shown to bind PAR. A previously characterized non-functional RNF146 WWE mutant variant (RFP-WWE mut., Andrabi *et al*., 2011) and free RFP served as negative controls. Asterisks indicate the expected molecular weights of the RFP-tagged proteins and the AtPARP2-GFP fusion protein. CBB, Coomassie Brilliant Blue stained PVDF membrane. The result is representative of three independent experiments.

### Ta-sro1 forms oligomers via the WWE-PARP domains

The N-terminal 265 amino acids of Arabidopsis RCD1 are sufficient for self-association in an Y2H assay. However, if the PARP and RST domains are included in one of the two Y2H constructs the interaction is strongly attenuated (Wirthmueller *et al*., 2018). Therefore, from this previous report it remains unclear if full-length SRO proteins form dimers or oligomers. During purification of recombinantly expressed Ta-sro1 we noticed that the 66 kDa protein eluted in two peaks from a preparative SEC column (Figs. S1, S2). The first peak corresponded to an apparent oligomeric complex of ∼266 kDa and when analyzed by SDS-PAGE resolved into two to three bands in the range of 55-70 kDa. The second peak (∼41 kDa) eluted closer to the expected molecular weight of monomeric Ta-sro1 and consisted of a 50 kDa band and several smaller bands (Fig. S2). An anti-His_6_ immunoblot indicated degradation of the protein from the C-terminus (Fig. S2). As we previously found evidence for SRO protein oligomerization via their N-terminal domain(s), we expressed and purified the Ta-sro1 WWE-PARP domains (amino acids 2-434) with a cleavable, N-terminal His_6_-tag. This truncated 50 kDa Ta-sro1 fragment showed no signs of degradation and eluted in a defined peak (∼206 kDa) in SEC on a preparative column after cleavage of the His_6_-tag (Fig. S2). On an analytical S200 SEC column the Ta-sro1 WWE-PARP construct eluted with an apparent molecular weight of 203 kDa and therefore earlier than the 150 kDa ADH and 66 kDa

BSA marker proteins (Fig. 5a). Addition of 10 mM DTT to the protein and the chromatography buffer only had a marginal effect on the elution profile. High ionic strength buffer conditions (0.5 M NaCl added to the protein sample and the buffer) resulted in an even smaller elution volume (Fig. 5a). SDS-PAGE of the Ta-sro1 WWE-PARP oligomer under reducing and non-reducing conditions revealed that the majority of the protein migrates as a monomer under non-reducing conditions (Fig. 5b). However, in comparison to reducing conditions we noticed two bands of higher molecular weight that could represent a disulfide-linked oligomeric complex. Therefore, whilst disulfides might contribute to Ta-sro1 oligomerization *in vitro* they are clearly not the major type of intermolecular interaction for the recombinantly expressed protein. We then tested if Ta-sro1 forms oligomers in plant cells. As shown in Fig. 5c, transiently expressed Ta-sro1-GFP protein co-immunoprecipitated HA-tagged Ta-sro1. Free GFP served as negative control. As the protein extraction and wash buffers contained 5 mM DTT, oligomerization in plant cells is probably not predominantly mediated by disulfides, a result that is consistent with our *in vitro* analyses. Although we did not detect a strong association between PAR chains and Ta-sro1 in plant cells, we tested if PAR affects the oligomeric state of the protein. We incubated the Ta-sro1 WWE-PARP domains with PAR (approximately 1:2 molar ratio with respect to the iso-ADP-ribose moiety) and repeated the analytical SEC assay. As shown in Fig. S3, the elution volume of the Ta-sro1 WWE-PARP protein was unaltered in presence of PAR. The A_280_/A_260_ ratio of the protein peak was similar in presence and absence of PAR indicating that at least the oligomeric form of the protein does not bind detectable amounts of PAR *in vitro*.

**Figure 5.**
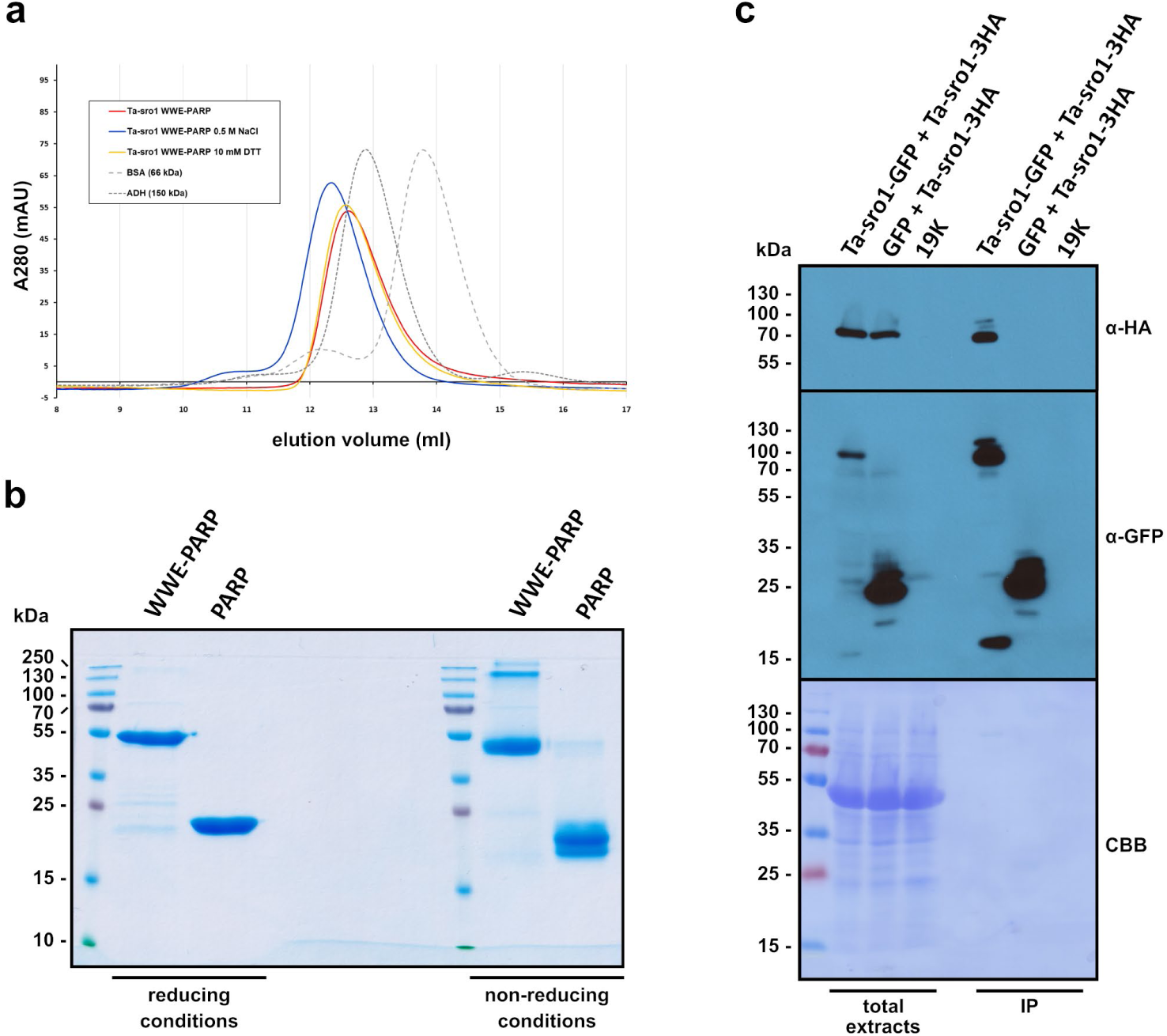
The Ta-sro1 WWE-PARP domains mediate formation of protein oligomers. **(a)** Elution profiles of the Ta-sro1 WWE-PARP domains from an analytical S200 size exclusion chromatography column under different buffer conditions. The protein was either analyzed in standard buffer (50 mM HEPES-NaOH pH 7.5, 150 mM NaCl) (red) or the same buffer supplemented with 10 mM DTT (yellow) or NaCl to a final concentration of 0.5 M (blue). Dashed lines indicate the elution profiles of the 150 and 66 kDa molecular weight marker proteins. **(b)** Recombinantly expressed and purified Ta-sro1 PARP domain or WWE-PARP domains were boiled in SDS-sample buffer with or without 100 mM DTT and separated by PAGE. **(c)** GFP- and 3HA- tagged variants of Ta-sro1 were co-expressed in *N. benthamiana.* A co-infiltration of Ta-sro1-3HA with GFP served as negative control. GFP and Ta-sro1-GFP were immunoprecipitated using magnetic GFP affinity beads and total extracts and immunoprecipitates were analyzed by α-HA and α-GFP immunoblots. CBB, Coomassie Brilliant Blue stained PVDF membrane. The result is representative of three independent experiments.

## Discussion

Our reassessment of the proposed Ta-sro1 PARP activity revealed that the previously reported catalytic activity could not be reproduced under our conditions and with appropriate controls (Fig. 3a). Furthermore, several molecular features that are eminent from the Ta-sro1 PARP domain structure are inconsistent with PARP activity. The His of the catalytic His-Tyr-Glu triad is replaced by Leu^312^ in Ta-sro1. Strict conservation of a His residue at this position in active ADP-ribosyltransferases is explained by its function in forming a hydrogen bond to the 2’-OH of the proximal ribose (Ruf *et al*., 1996; Steffen *et al*., 2013; Langelier *et al*., 2018). Consequently, this conserved His is essential for HsPARP1 activity (Marsischky *et al*., 1995; Shao *et al*., 2020) and a corresponding His^21^ to Leu exchange in the structurally related diphtheria toxin abolishes ADP-ribosylation (Johnson & Nicholls, 1994; Bell & Eisenberg, 1997). In active ADP-ribosyltransferases the His is followed by a conserved Gly residue that forms two hydrogen bonds with the amide group of NAD^+^ (Wahlberg *et al*., 2012; Langelier *et al*., 2018). As shown in Fig. 1d, Ta-sro1 Pro^313^ at this position would sterically interfere with accommodation of the nicotinamide moiety and consistently a corresponding Gly to Trp mutation in HsPARP10 abolishes ADP-ribosyltransferase activity (Yu *et al*., 2005). Finally, the Ta-sro1 PARP domain lacks an extended acceptor loop that appears to function in binding target proteins and/or the elongating PAR chain in canonical PARPs (Ruf *et al*., 1998; Kleine *et al*., 2008; Vyas *et al*., 2014). These results are conflicting with the previously proposed model claiming that elevated PARP activity of the hypermorphic Ta-sro1 proteoform is responsible for the higher salinity stress tolerance of wheat cultivar SR3 (Liu *et al*., 2014).

The Ta-SRO1 gene from the parental wheat cultivar JN177 differs in three nucleotides from Ta-sro1 (Liu *et al*., 2014). Two of these nucleotide polymorphisms alter the amino acid sequence resulting in the hypermorphic Ta-sro1 proteoform. Liu *et al*. (2014) suggested that the three nucleotide polymorphisms are a consequence of genomic stress induced during the asymmetric genome hybridization between JN177 and *T. ponticum*. However, subsequently released sequences of wheat 5B chromosomes revealed that predicted coding sequences with 100% identity exist in durum wheat and in bread wheat cultivar Chinese Spring (International Wheat Genome Sequencing Consortium (IWGSC) *et al*., 2018; NCBI Genbank entries KAF7069564.1 and VAI40608.1, Fig. S4a). Therefore, recombination with genetic material from another cultivar is a more plausible scenario for introduction of the Ta-sro1 allele into cultivar SR3. A Thr at the position that corresponds to Ta-sro1 Thr^343^ is also present in other SRO proteins from several monocot species (Fig. S4b). Some of these homologs also have a Val at the position corresponding to Ta-sro1 Val^250^. Therefore, the Val^250^/Thr^343^ combination is not unique to the wheat Ta-sro1 proteoform.

Our Ta-sro1 PARP domain structure pinpoints the location of the two polymorphic amino acids (Fig. 1b) but functional consequences of the amino acid exchanges remain to be determined. The position of the Gly/Val^250^ polymorphism at the very N-terminus of the crystallized construct complicates interpretations with respect to Ta-sro1 function. We noticed that the NESmapper algorithm for predicting putative nuclear export signals (NES) identifies the Ta-sro1 peptide GQPVDSAVRKLLLE (247-260) as a likely NES with a score of 15.1 (Kosugi *et al*., 2014). The exchange of Val^250^ to Gly lowers the NESmapper score to 2.5 indicating that the two proteoforms could have different nuclear export rates. Active nuclear export of SRO proteins is not unprecedented as a putative NES in Arabidopsis RCD1 has been identified (Shapiguzov *et al*., 2019) and a predominantly cytoplasmic localization of RCD1 has been observed under high salinity stress (Katiyar-Agarwal *et al*., 2006). However, when we expressed GFP-tagged variants of the two Ta-SRO1 proteoforms in *N. benthamiana,* both proteins appeared entirely nuclear localized suggesting that at least steady-state transport kinetics over the nuclear envelope are not substantially different (Fig. S5). The putative Ta-sro1 NES residues form a short α-helix that connects to the core of the PARP domain by the Gly^261^-Ala-Ala-Gly^264^ linker. This N-terminal α-helix associates with the core PARP domain by hydrophobic interactions and by two salt bridges formed between Asp^251^/Arg^276^ and Arg^255^/Glu^268^, respectively. Therefore, accessibility of the putative NES for binding to nuclear transport factors would require a substantial conformational change at the N-terminal region of the PARP domain.

The second polymorphic residue (Ala/Thr^343^) maps to the proposed active site of the Ta-sro1 PARP domain (Fig. 1b, c). In human PARP1 the corresponding amino acid has been identified as a ’gatekeeper’ residue that, when mutated, widens the NAD^+^ binding site for access to bulkier NAD^+^ derivatives (Gibson *et al*., 2016). Based on protein homology modeling, Liu *et al*. (2014) proposed that the Ala/Thr^343^ side chain interacts with the co-substrate NAD^+^. However, our Ta-sro1 PARP domain crystal structure reveals that the Ta-sro1 Thr^343^ side chain points in the opposite direction and towards a hydrophobic area forming the basis of the postulated NAD^+^ binding site (Fig. 1c). Conceivably, exchange of a hydrophobic to a polar side chain at this position could slightly alter the position of strand β3 that forms one side of the pocket. This might affect a so far uncharacterized enzymatic activity of Ta-SRO1. However, at least for Arabidopsis RCD1, several amino acid exchanges at the presumed active site do not compromise the function of the protein in plants (Wirthmueller *et al*., 2018).

Why PARP domains are strictly conserved in SRO proteins if they do not show ADP-ribosyl-transferase activity, remains an outstanding question. Several studies indicate an alternative role of SRO PARP domains in binding to transcription factors, presumably in cooperation with the C-terminal RST domain. The PARP domain of *Musa acuminata* SRO4 is required for binding to the transcription factor MaMYB4 (Zhang *et al*., 2019). In rice, the OsSRO1c PARP domain appears to be the predominant interaction site for several transcription factors (You *et al*., 2013, 2014). Complex formation between Arabidopsis RCD1 and transcription factors was first identified with an RCD1 fragment that includes the PARP and RST domains (Jaspers *et al*., 2009). Subsequent analyses showed that several transcription factors bind to RCD1 via short linear motifs and that the isolated RST domain is sufficient to mediate these interactions with dissociation constants in the sub-micromolar range (Kjaersgaard *et al*., 2011; Vainonen *et al*., 2012; O’Shea *et al*., 2015; Bugge *et al*., 2018; Christensen *et al*., 2019). Therefore, the C-terminal RST domain is the primary site of interaction between transcription factors and SRO proteins but it remains possible that the PARP domain contributes to, or mediates, specific types of interactions.

The recent finding that the RCD1 WWE domain is a PAR chain-binding module by Vainonen *et al*. (2021) suggests that plant WWE domains can retain the ability to recognize iso-ADP-ribose moieties despite substantial sequence divergence from their mammalian homologs (Wang *et al*., 2012). Here, we demonstrate that at least in plant cell extracts, the Ta-sro1 protein does not form detectable protein complexes with an auto-PARylated bait protein (Fig. 4). Similarly, the apparent WWE-PARP domain oligomer does not bind PAR in SEC (Fig. S5). Our attempts to address Ta-sro1 PAR binding *in vitro* using a dot blot assay were inconsistent and further quantitative binding studies are necessary to probe for this association *in vitro*. Nevertheless, we can infer from the *in planta* PAR chain binding assay that Ta-sro1 shows a substantially lower affinity to PAR chains compared to the previously characterized RNF146 WWE domain (Zhang *et al*., 2011; Wang *et al*., 2012). An additional or alternative function of the WWE-IDR2 region is the formation of SRO oligomers. From previous analyses and results presented here, it appears that SRO proteins form at least two different types of higher-order protein complexes. A disulfide-linked oligomer of Arabidopsis RCD1 can be specifically induced by methyl viologen (MV) or H_2_O_2_ treatment (Shapiguzov *et al*., 2019). However, also under mildly reducing conditions, the RCD1 WWE-IDR2 region immunoprecipitates endogenous RCD1 from non-stressed Arabidopsis protein extracts (Wirthmueller *et al*., 2018). Although based on over-expression of the isolated WWE-IDR2 protein fragment, this result is indicative of a pre-formed RCD1 oligomer that may serve as pre-assembly state for ROS-induced disulfide-linked protein complexes (Shapiguzov *et al*., 2019). Here, we demonstrate that DTT-insensitive oligomers of full-length Ta-sro1 exist in plant cell extracts and that the recombinantly expressed protein forms a similar higher order complex *in vitro*. This suggests that oligomer formation is an inherent property of WWE-PARP-RST (type A) SRO proteins and may have implications for the function of type B SROs that lack the WWE-IDR2 region (Jaspers *et al*., 2010).

An altered redox homeostasis is a common consequence of mutations in SRO genes. Wheat cultivar SR3 shows a higher ROS (H_2_O_2_) accumulation under stress but also under control conditions (Liu *et al*., 2014). Moreover, overexpression of the Ta-sro1 proteoform in Arabidopsis renders the transgenic plants more tolerant to MV, a herbicide that accepts electrons from photosystem I and transfers them to molecular oxygen thereby leading to ROS accumulation in chloroplasts (Liu *et al*., 2014). Similar results have been reported for overexpression of the maize *SRO1b* gene in Arabidopsis. *ZmSRO1b* overexpression not only confers higher MV tolerance but also mitigates the effects of high salinity on plant growth (Li *et al*., 2018). Therefore, overexpression of Ta-sro1 or *ZmSRO1b* phenotypically mimic the elevated MV tolerance of Arabidopsis *rcd1* mutants (Ahlfors *et al*., 2004). Likewise, rice sro1c-1 knock-out mutant are less sensitive to MV (You *et al*., 2013). These results show that an altered redox homeostasis can be caused by loss of specific SRO isoforms, ectopic overexpression of SRO genes from other plant species, or expression of specific SRO proteoforms. Given the role of ROS in early signal transduction and cell-to-cell communication (Gilroy *et al*., 2014; Evans *et al*., 2016), the constitutively elevated ROS levels in wheat cultivar SR3 may underlie the enhanced salinity tolerance phenotype (Liu *et al*., 2014). However, the results presented here argue against the current hypothesis that elevated PARP activity of the hypermorphic Ta-sro1 proteoform is the molecular basis for the enhanced SR3 salinity tolerance. Mapping the two polymorphic amino acids onto the PARP domain structure provides a structural basis for exploring differences between the two proteoforms in the future. However, our results also suggest that such a functional characterization of the two proteoforms will require a better understanding of how Ta-SRO1 functions as a transcriptional co-regulator and that this likely involves inter- and intra-protein interactions.

## Acknowledgments

We thank Claudia Alings, Freie Universität Berlin, for technical assistance with protein crystallization. We acknowledge access to beamlines of the BESSY II storage ring (Berlin, Germany) via the Joint Berlin MX-Laboratory sponsored by the Helmholtz Zentrum Berlin für Materialien und Energie, the Freie Universität Berlin, the Humboldt-Universität zu Berlin, the Max-Delbrück-Centrum, the Leibniz-Institut für Molekulare Pharmakologie and Charité – Universitätsmedizin Berlin. LW acknowledges funding by the German Research Foundation (DFG grant WI 3670/2-1) and core funding from the Leibniz Institute of Plant Biochemistry (IPB) and the Freie Universität Berlin Dahlem Centre of Plant Sciences.

## Author contributions

KF, SH, RDM, BL, and LW conceived and designed experiments. SV, KF, SH, RDM, RL, BL, and LW carried out experiments. SV, KF, SH, BL, and LW analyzed the data. KF, BL, and LW wrote the manuscript. All authors reviewed and approved the submitted manuscript.

---

**Supplementary Figure S1.**
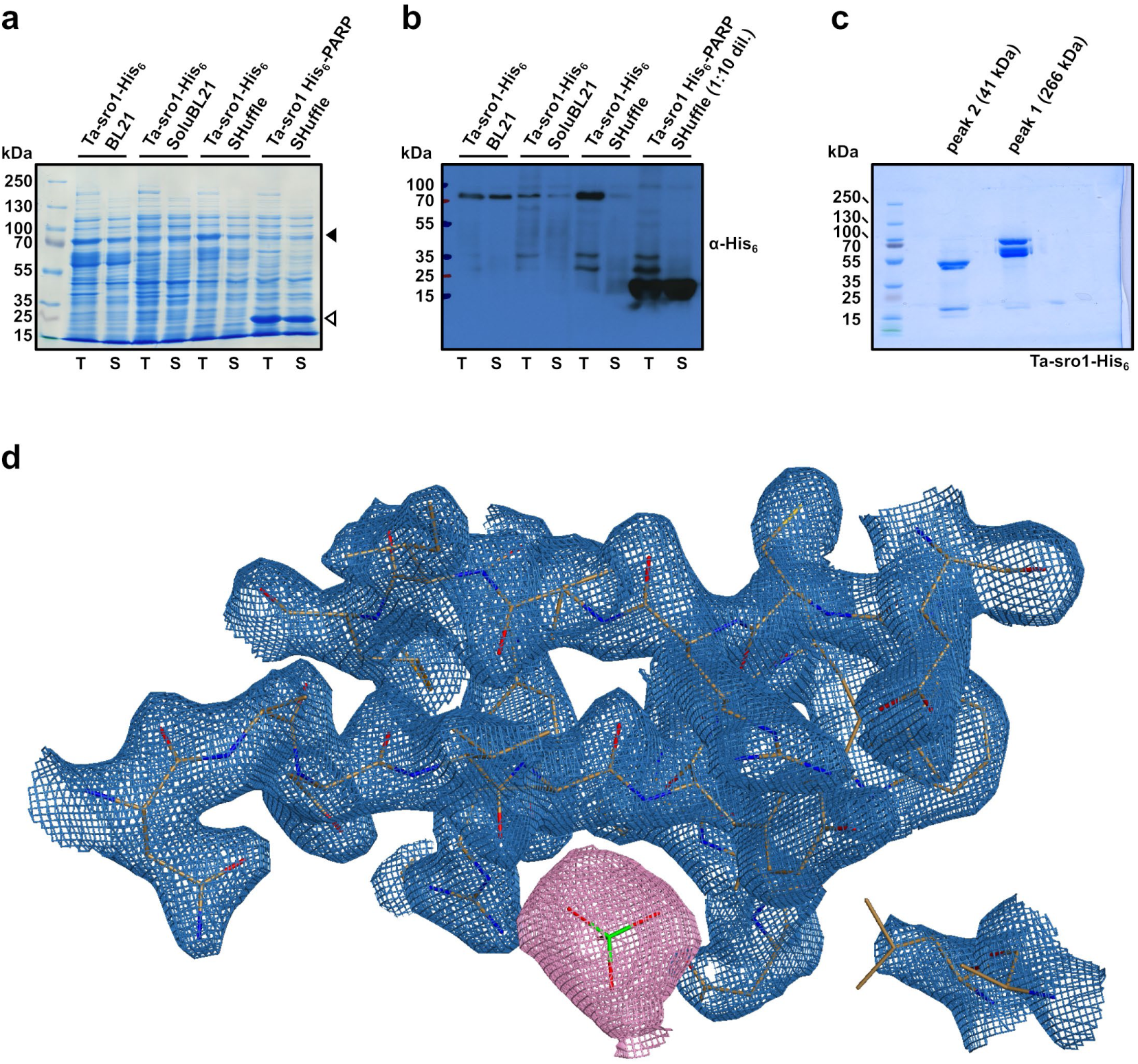
Recombinant expression and purification of Ta-sro1-His_6_ and the His_6_- tagged PARP domain. **(a)** Total (T) and soluble (S) fractions of PARP domain expressed in *E. coli* Shuffle cells and of the full-length Ta-sro1-His_6_ protein expressed in different *E. coli* strains. Closed and open arrowheads indicate migration of the Ta-sro1-His_6_ protein and the His_6_-tagged PARP domain, respectively. **(b)** α-His_6_ immunoblot of the samples shown in (a). **(c)** SDS-PAGE of the two Ta-sro1-His_6_ elution peaks from a preparative S200 size exclusion chromatography column. **(d)** Polder electron density maps (Liebschner *et al*., 2017) shown as pink mesh at a σ-level of 3.0 around the sulfate molecule, omitted for map calculation. Blue mesh, 2Fo-Fc electron density for the surrounding protein environment. Amino acid residues are shown in stick representation.

**Supplementary Figure S2.**
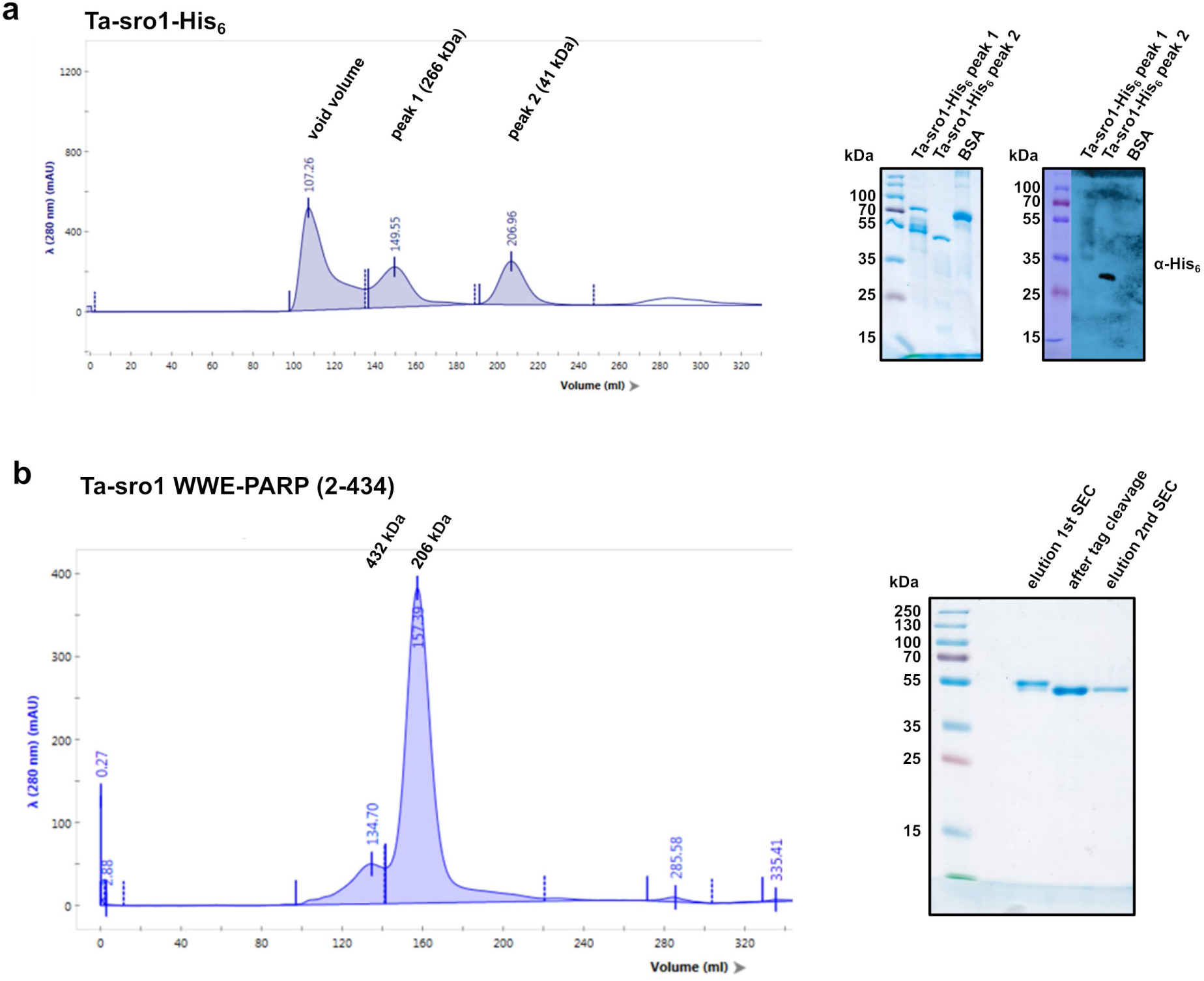
Elution profiles of Ta-sro1-His_6_ and the purified WWE-PARP domains in size exclusion chromatography. **(a)** Elution profile of Ta-sro1-His_6_ protein from a preparative S200 size exclusion chromatography column and analysis of the two indicated protein peaks by SDS-PAGE and α-His_6_ immunoblot. **(b)** Elution profile of the purified Ta-sro1 WWE-PARP domains after cleavage of the His_6_-tag and SDS-PAGE of the protein before and after tag cleavage.

**Supplementary Figure S3.**
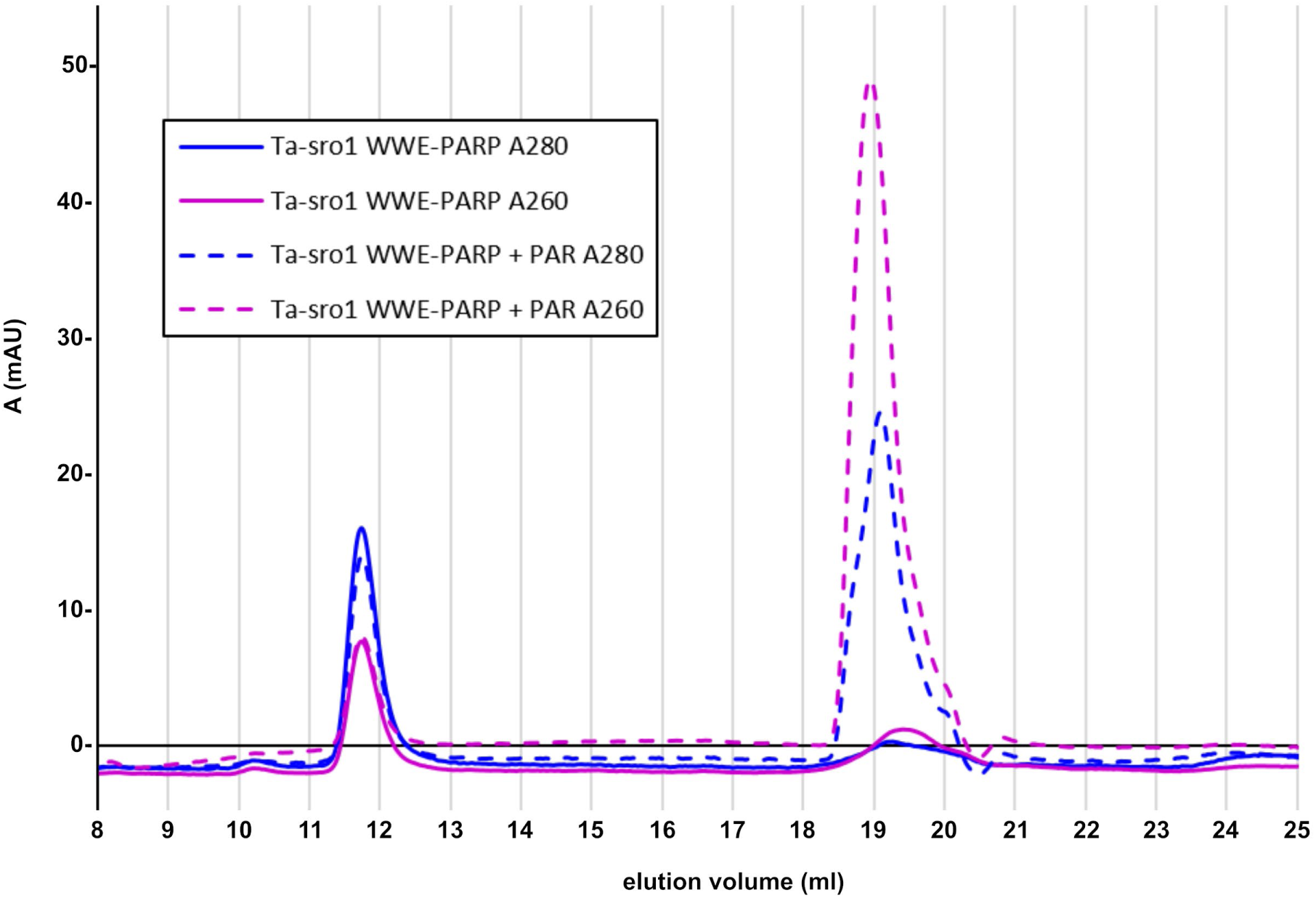
Elution profiles of the Ta-sro1 WWE-PARP domains in absence and presence of PAR. Ta-sro1 WWE-PARP protein at a final concentration of 1.43 μM was incubated in presence or absence of 4.29 μM PAR (approximately 1:2 molar ratio with respect to the iso-ADP-ribose moiety in PAR) for 30 min at 20 °C. Subsequently, the protein was analyzed on an analytical S200 size exclusion chromatography column. Solid curves show the A_260_ and A_280_ nm absorption spectra of the protein without PAR. Dashed curves denote the absorption spectra in presence of PAR. Protein and PAR can be distinguished by the higher A_260_/A_280_ ratio of PAR.

**Supplementary Figure S4.**
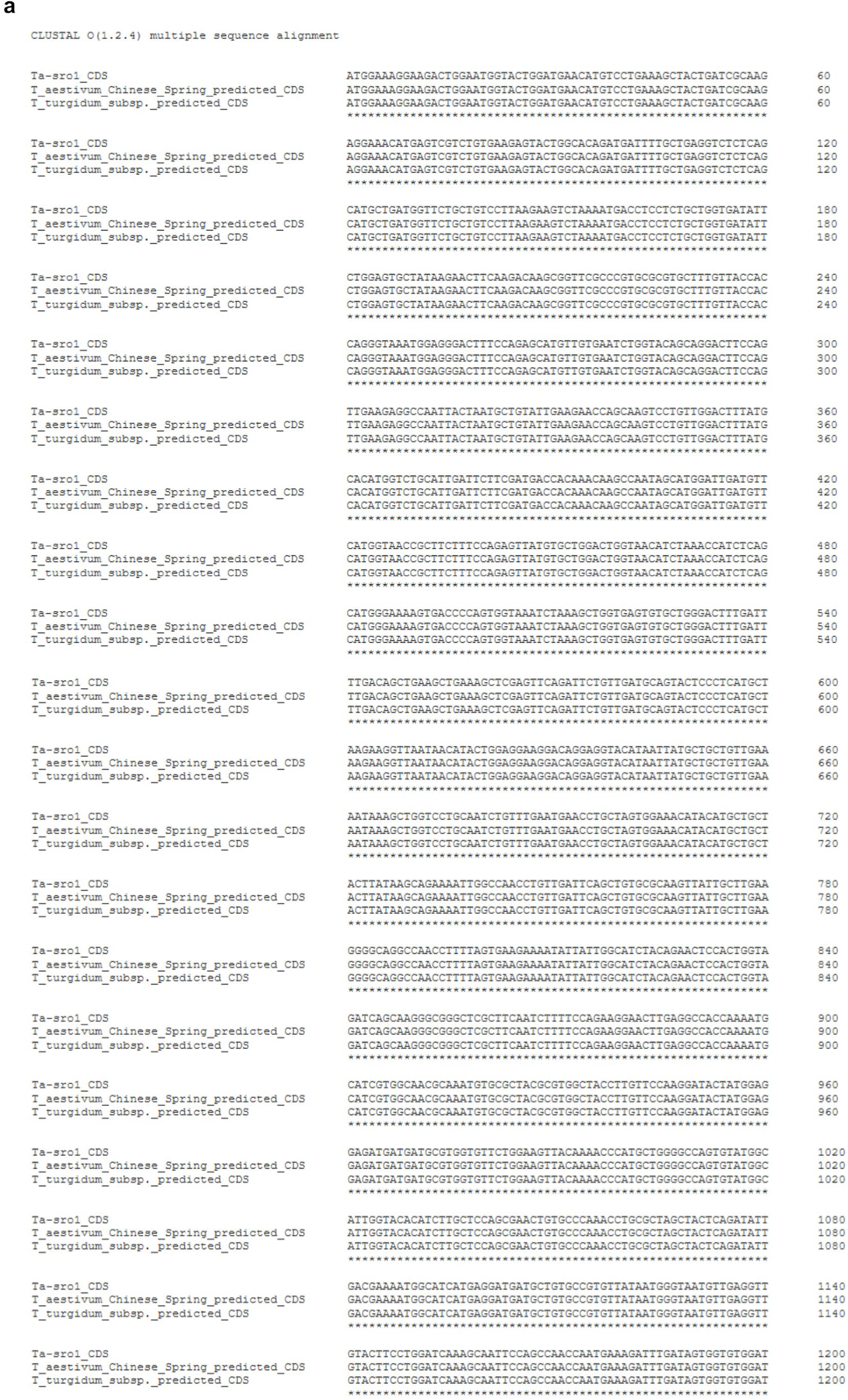
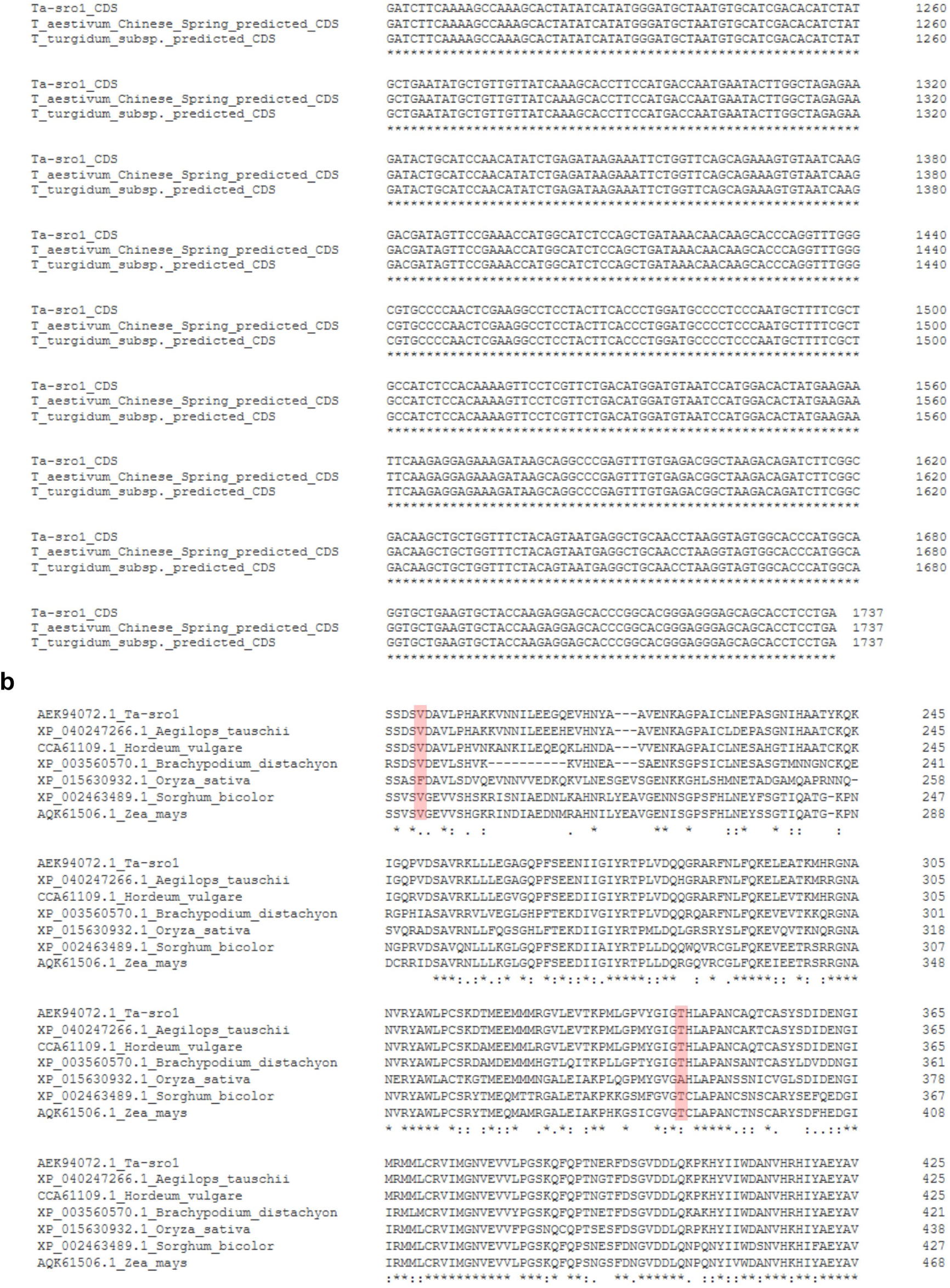
The Ta-sro1 Gly^250^ to Val and Ala^343^ to Thr polymorphisms are not unique to Ta-sro1. **(a)** Coding sequences with 100% identity to Ta-sro1 identified via the NCBI ’identical proteins’ search option. The coding sequences of wheat cultivar Chinese Spring and *Triticum turgidum* subsp. *durum* correspond to NCBI Genbank entries KAF7069564.1 and VAI40608.1, respectively. **(b)** Val and Thr residues in positions that correspond to Ta-sro1 Val^250^ and Thr^343^ are present in SRO proteins from different monocot species. Nucleotide and protein sequences were aligned with Clustal Omega (Sievers *et al*., 2011).

**Supplementary Figure S5.**
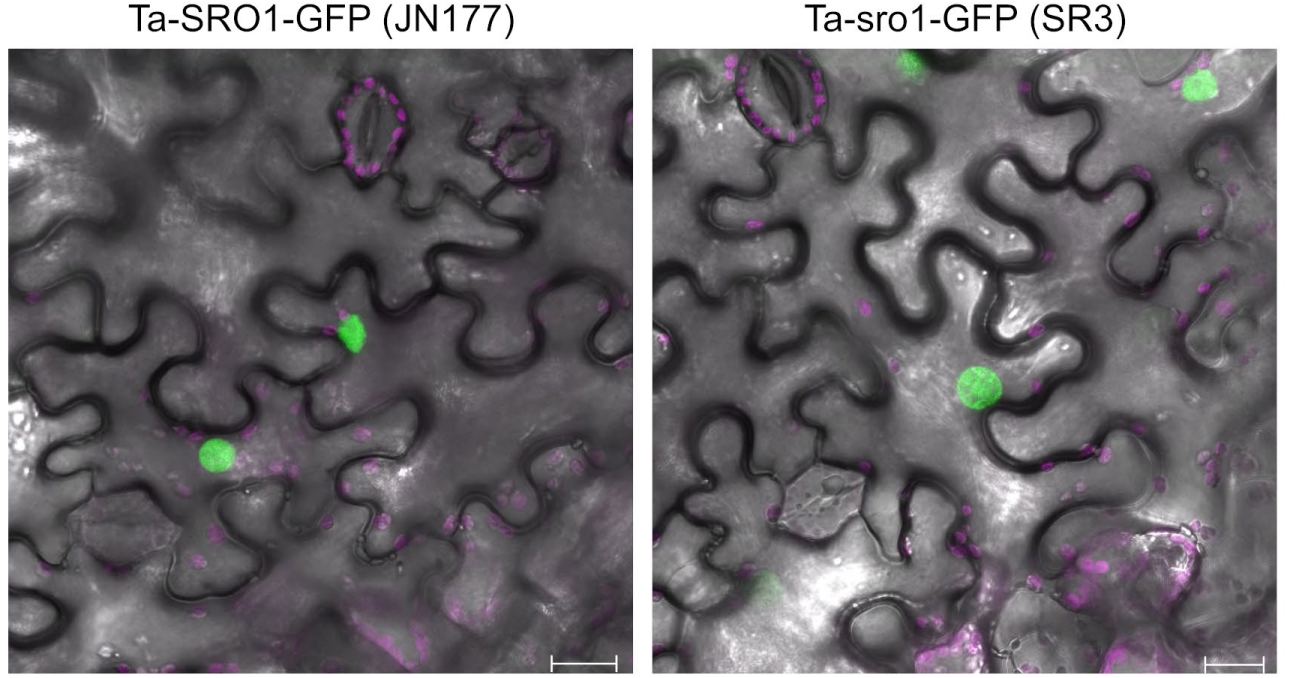
The Ta-sro1 (SR3) and Ta-SRO1 (JN177) proteoforms do not show differences in subcellular localization when transiently expressed in *N. benthamiana*. GFP-tagged versions of the two proteoforms were analyzed by confocal microscopy. The GFP signal is false-colored green, purple color shows chlorophyll fluorescence from chloroplasts. Scale bars = 20 μm. The images are representative of five independent leaf sections per proteoform monitored in a single experiment.

**Supplementary Table S1.**
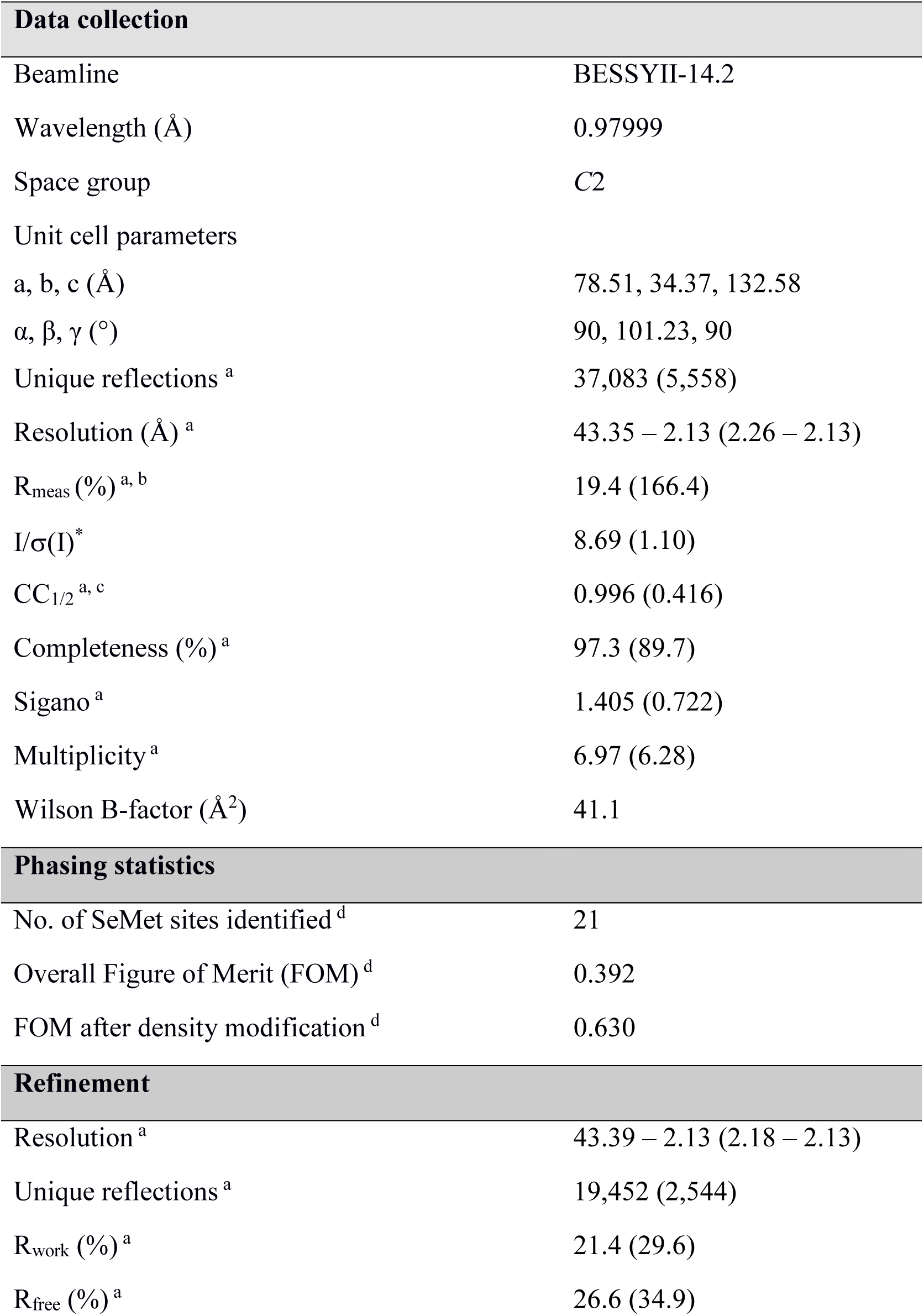
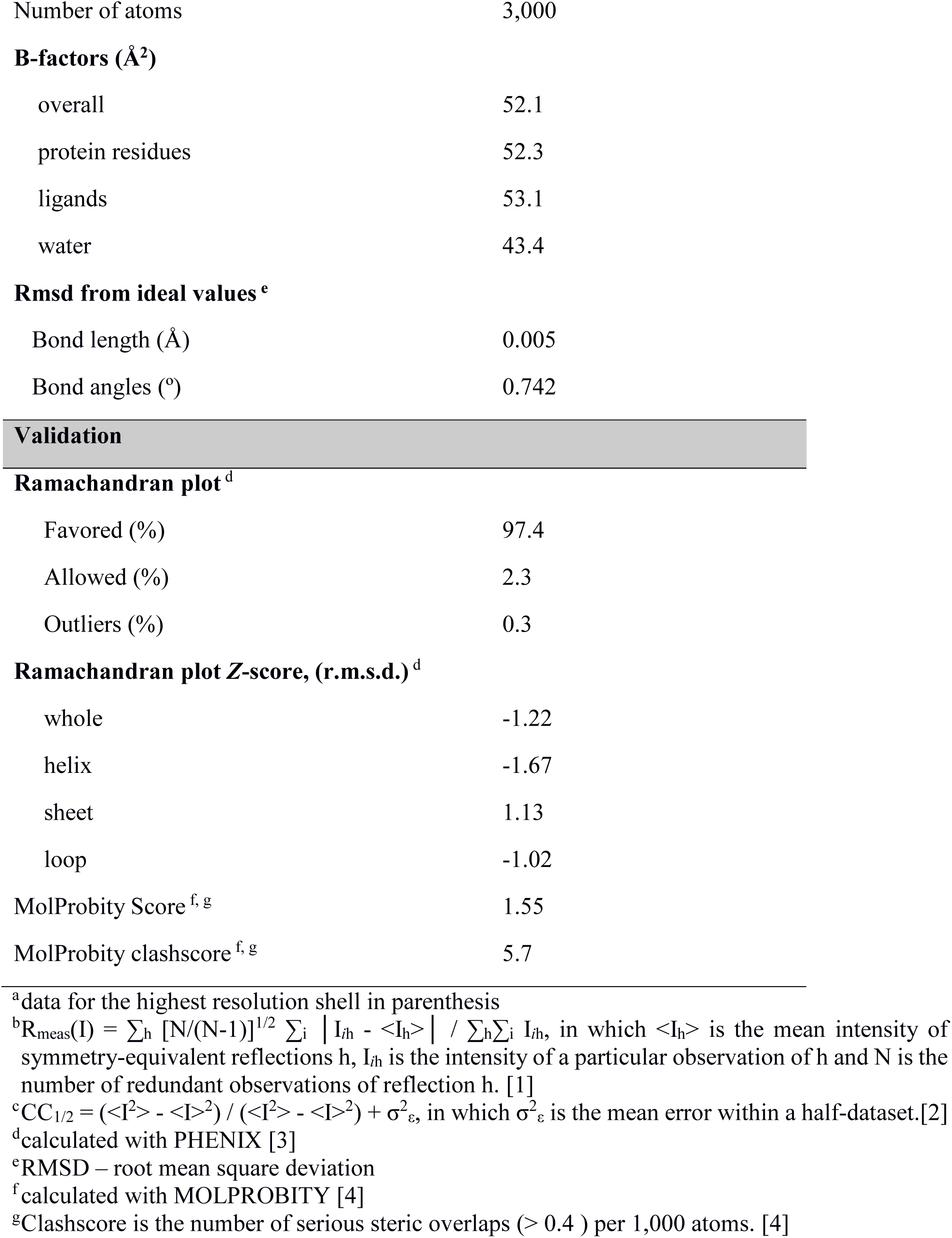
X-ray data collection, refinement, and validation statistics of the Ta-sro1 PARP domain structure.

| **Data collection** |  |
| --- | --- |
| Beamline | BESSYII-14.2 |
| Wavelength (Å) | 0.97999 |
| Space group | *C2* |
| Unit cell parameters |  |
| a, b, c (Å) | 78.51, 34.37, 132.58 |
| $\alpha$, $\beta$, $\gamma$ (°) | 90, 101.23, 90 |
| Unique reflections <sup>a</sup> | 37,083 (5,558) |
| Resolution (Å) <sup>a</sup> | 43.35 – 2.13 (2.26 – 2.13) |
| R<sub>meas</sub> (%) <sup>a, b</sup> | 19.4 (166.4) |
| I/$\sigma$(I) <sup>*</sup> | 8.69 (1.10) |
| CC<sub>1/2</sub> <sup>a, c</sup> | 0.996 (0.416) |
| Completeness (%) <sup>a</sup> | 97.3 (89.7) |
| Sigano <sup>a</sup> | 1.405 (0.722) |
| Multiplicity <sup>a</sup> | 6.97 (6.28) |
| Wilson B-factor (Å<sup>2</sup>) | 41.1 |
| **Phasing statistics** |  |
| No. of SeMet sites identified <sup>d</sup> | 21 |
| Overall Figure of Merit (FOM) <sup>d</sup> | 0.392 |
| FOM after density modification <sup>d</sup> | 0.630 |
| **Refinement** |  |
| Resolution <sup>a</sup> | 43.39 – 2.13 (2.18 – 2.13) |
| Unique reflections <sup>a</sup> | 19,452 (2,544) |
| R<sub>work</sub> (%) <sup>a</sup> | 21.4 (29.6) |
| R<sub>free</sub> (%) <sup>a</sup> | 26.6 (34.9) |
| Number of atoms | 3,000 |
| **B-factors (Å<sup>2</sup>)** |  |
| overall | 52.1 |
| protein residues | 52.3 |
| ligands | 53.1 |
| water | 43.4 |
| **Rmsd from ideal values <sup>e</sup>** |  |
| Bond length (Å) | 0.005 |
| Bond angles (°) | 0.742 |
| **Ramachandran plot <sup>d</sup>** |  |
| Favored (%) | 97.4 |
| Allowed (%) | 2.3 |
| Outliers (%) | 0.3 |
| **Ramachandran plot Z-score, (r.m.s.d.) <sup>d</sup>** |  |
| whole | -1.22 |
| helix | -1.67 |
| sheet | 1.13 |
| loop | -1.02 |
| MolProbity Score <sup>f, g</sup> | 1.55 |
| MolProbity clashscore <sup>f, g</sup> | 5.7 |

<sup>a</sup>data for the highest resolution shell in parenthesis
<sup>b</sup> $R_{\text{meas}}(I) = \sum_h [N/(N-1)]^{1/2} \sum_i |I_{ih} - \langle I_h \rangle| / \sum_h \sum_i I_{ih}$ , in which $\langle I_h \rangle$ is the mean intensity of symmetry-equivalent reflections $h$ , $I_{ih}$ is the intensity of a particular observation of $h$ and $N$ is the number of redundant observations of reflection $h$ . [1]
<sup>c</sup> $CC_{1/2} = (\langle I^2 \rangle - \langle I \rangle^2) / (\langle I^2 \rangle - \langle I \rangle^2) + \sigma_{\epsilon}^2$ , in which $\sigma_{\epsilon}^2$ is the mean error within a half-dataset.[2]
<sup>d</sup>calculated with PHENIX [3]
<sup>e</sup>RMSD – root mean square deviation
<sup>f</sup>calculated with MOLPROBITY [4]
<sup>g</sup>Clashscore is the number of serious steric overlaps ( $> 0.4$ ) per 1,000 atoms. [4]

**Supplementary Table S2.** List of oligonucleotide sequences.

